# Inferring brain-wide interactions using data-constrained recurrent neural network models

**DOI:** 10.1101/2020.12.18.423348

**Authors:** Matthew G. Perich, Charlotte Arlt, Sofia Soares, Megan E. Young, Clayton P. Mosher, Juri Minxha, Eugene Carter, Ueli Rutishauser, Peter H. Rudebeck, Christopher D. Harvey, Kanaka Rajan

**Affiliations:** Icahn School of Medicine at Mount Sinai, New York, NY, USA; Harvard Medical School, Boston, MA, USA; Cedars-Sinai Medical Center, Los Angeles, CA, USA; California Institute of Technology, Pasadena, CA

## Abstract

Behavior arises from the coordinated activity of numerous anatomically and functionally distinct brain regions. Modern experimental tools allow unprecedented access to large neural populations spanning many interacting regions brain-wide. Yet, understanding such large-scale datasets necessitates both scalable computational models to extract meaningful features of inter-region communication and principled theories to interpret those features. Here, we introduce Current-Based Decomposition (CURBD), an approach for inferring brain-wide interactions using data-constrained recurrent neural network models that directly reproduce experimentally-obtained neural data. CURBD leverages the functional interactions inferred by such models to reveal directional currents between multiple brain regions. We first show that CURBD accurately isolates inter-region currents in simulated networks with known dynamics. We then apply CURBD to multi-region neural recordings obtained from mice during running, macaques during Pavlovian conditioning, and humans during memory retrieval to demonstrate the widespread applicability of CURBD to untangle brain-wide interactions underlying behavior from a variety of neural datasets.

## INTRODUCTION

During development, the nervous systems of even small organisms organize into remarkably complex structures. Brains exhibit structural modularity (e.g., brain regions, laminar organization, cell types) with phylogenetically-determined specialization across modules^1^. Brain regions, in particular, have striking specialization and unique functional characteristics. However, individual brain regions also frequently interact with numerous other regions throughout the brain^2^. These macroscopic circuits are recurrently connected via direct projections, multi-synapse loops, and more widespread, indirect effects such as neuromodulator release^3^. Consequently, much of the brain is active during even simple behaviors that could, in theory, be mediated by only a smaller subset of regions^4–6^. Deriving an understanding of the neural basis of behavior requires consideration of the distributed nature of brain-wide activity. However, despite the prevalence of large-scale, multi-region datasets afforded by modern experimental techniques, researchers lack a comprehensive, unifying approach to infer brain-wide interactions and information flow. Here, we introduce Current-Based Decomposition (CURBD), a computational framework that leverages recurrent neural network (RNN) models of multi-region neural recordings to infer the magnitude and directionality of the interactions between regions across the brain. While most neural data analysis and dimensionality reduction techniques^7^ describe the output of neurons (e.g. spiking activity), CURBD reconceptualizes the activity of a neural population in terms of the inputs driving the neurons. We first introduce the conceptual advantages of CURBD and validate the method on simulated datasets where ground truth multi-region interactions are known. We then apply CURBD to multi-region calcium fluorescence recordings from four cortical regions of mice during running, and electrophysiological data from three cortical and subcortical regions in the rhesus macaque during Pavlovian conditioning and four brain regions of human participants during memory retrieval. These examples highlight the widespread applicability of CURBD for inferring multi-region interactions from large-scale neural datasets.

### Current-based decomposition of multi-region datasets using recurrent neural networks

CURBD operates on the fundamental premise that the exchange of currents between active units in a recurrently-connected neural network can be precisely estimated. In a single-layer network, the currents driving a single target unit can be viewed as a weighted sum of the activity of the “source” units (**Figure 1a**). Mathematically, these weights correspond to interaction strengths, summarized by a vector with each source unit represented as a single entry in the vector. However, neural circuits in biological brains are typically intricately, recurrently connected^2^. This feature prompted common use of RNNs to model their computational functions^8,9^. RNNs trained to produce desired behaviors^10–12^ and tasks^13–18^ or match neural data^2,17,19^ (or both^20,86^) can be reverse-engineered to generate hypotheses for how biological neural circuits could implement similar functions^21,22^. As in the single-layer network, the activity of any unit in an RNN can be computed as a weighted sum of the activity of all other units in the network, which are the sources of its input (**Figure 1b**). The activity of the network can thus be described compactly using a single “directed interaction” matrix quantifying the magnitude and type (excitatory or inhibitory) of the interactions.

**Figure 1.**
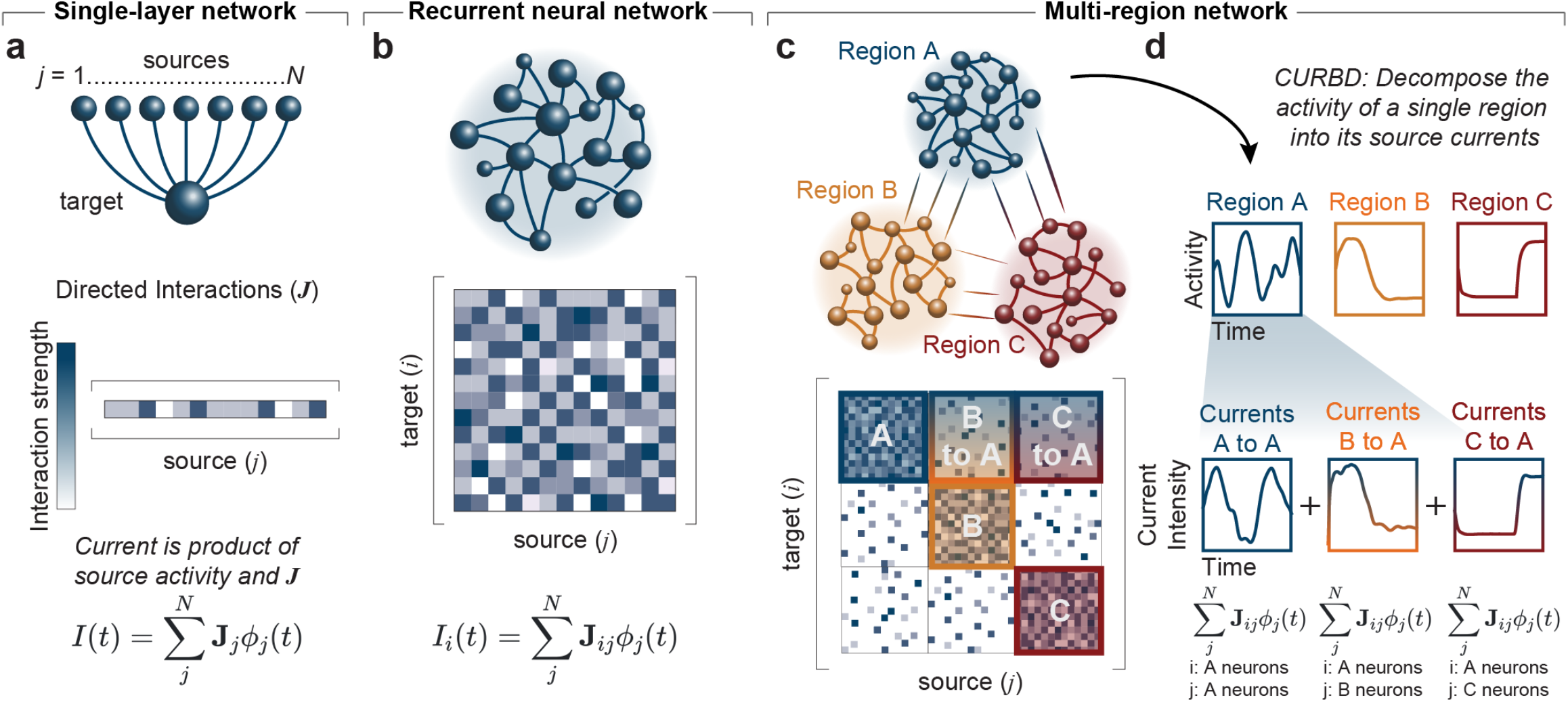
Current-based Decomposition (CURBD) of multi-region interactions using recurrent neural networks. **(a)** In a single-layer network, each source unit connects to a target unit with a directed interaction weight given by the vector *J*. The activity of the target unit can be derived based on its source current, a weighted combination of the source activity (*ϕ*) multiplied by the corresponding excitatory or inhibitory directed interaction weight. **(b)** In recurrent neural network (RNN) models, each unit is driven by inputs from the other units, but also sends outputs to those same units. Thus, the directed interactions are summarized by a matrix **J** with each column containing the weights of a source unit and each row those of a target unit. **(c)** Neural circuits can be modeled as a ‘network of networks’, i.e., with interconnected but distinct regions. The directed interactions governing multiple regions are still summarized by a single matrix **J** where submatrices along the diagonal correspond to within-region interactions, and off-diagonal submatrices correspond to interactions between different regions. **(d)** As in the single-layer network in Panel a and single-module RNN in Panel b, the currents driving each target unit can be viewed as the weighted sum of source activity from each of these submatrices. By multiplying the weights in each submatrix by the source activity of each region individually, we can decompose the total activity of any region (e.g., Region A) into the constituent source currents of the total activity.

Given the high degree of recurrent connectivity between regions, interactions between active neurons in different brain regions can be conceived as an RNN^2^. To implement CURBD, we model brain-wide circuitry as multiple inter-connected RNNs forming a “network of networks”^2^. The activity of units in each region of this RNN is shaped by excitatory and inhibitory “source currents” from all regions that provide input, including from recurrently connected units within the same region. If the connectivity relating these networks is known, then the source currents into a target region from any other region can be estimated using the corresponding submatrix of the directed interaction matrix and the activity of the source region (**Figure 1c**). When summed, these currents reconstruct the total activity of each neuron in the region, however, CURBD allows the total activity of each region to be decomposed into a set of source currents from all other regions (**Figure 1d**). Estimating population-wide inputs at this scale produces an unprecedented view into multi-region interactions. Furthermore, CURBD scales readily beyond two interconnected regions to brain-wide interactions^19^, circumventing a limitation of many existing approaches^23–26^.

### Implementation of CURBD

CURBD is based on the directed interaction matrix, **J**, which we use to infer currents. This matrix estimates the effective strength and type—excitatory (**J**_ij_>0) or inhibitory (**J**_ij_<0)— of interactions between active neurons, both within and across regions, that give rise to experimentally-observed neural dynamics. Since this matrix dictates the entire neural dynamical system over time, it captures the stability as well as the population-wide covariance of the activity (**Figure 2c**). Yet, a matrix capturing both the stability and structure of multi-region interactions is impossible to obtain through experimental measurements alone. Thus, we employ Model RNNs to infer the directed interaction matrix directly from multi-region experimental data obtained from behaving animals (**Figure 2a**). We first initialize a Model RNN with random connectivity. The Model RNN typically contains a number of units equal to the number of neurons available in the dataset to be modeled, but larger and subsampled variants can be employed^17^. Each model unit is then assigned to one neuron in the experimental dataset during training. The goal is to learn a directed interaction matrix such that the Model RNN autonomously reproduces the time-series activity of the recorded neurons given only the initial state of the neurons. At each time step, the activity of the next time point in the Model RNN is computed as the sum of the current state of population activity, *ϕ(t)*, weighted by **J** (Equation 1).

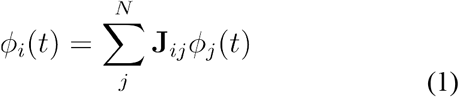

**Figure 2.**
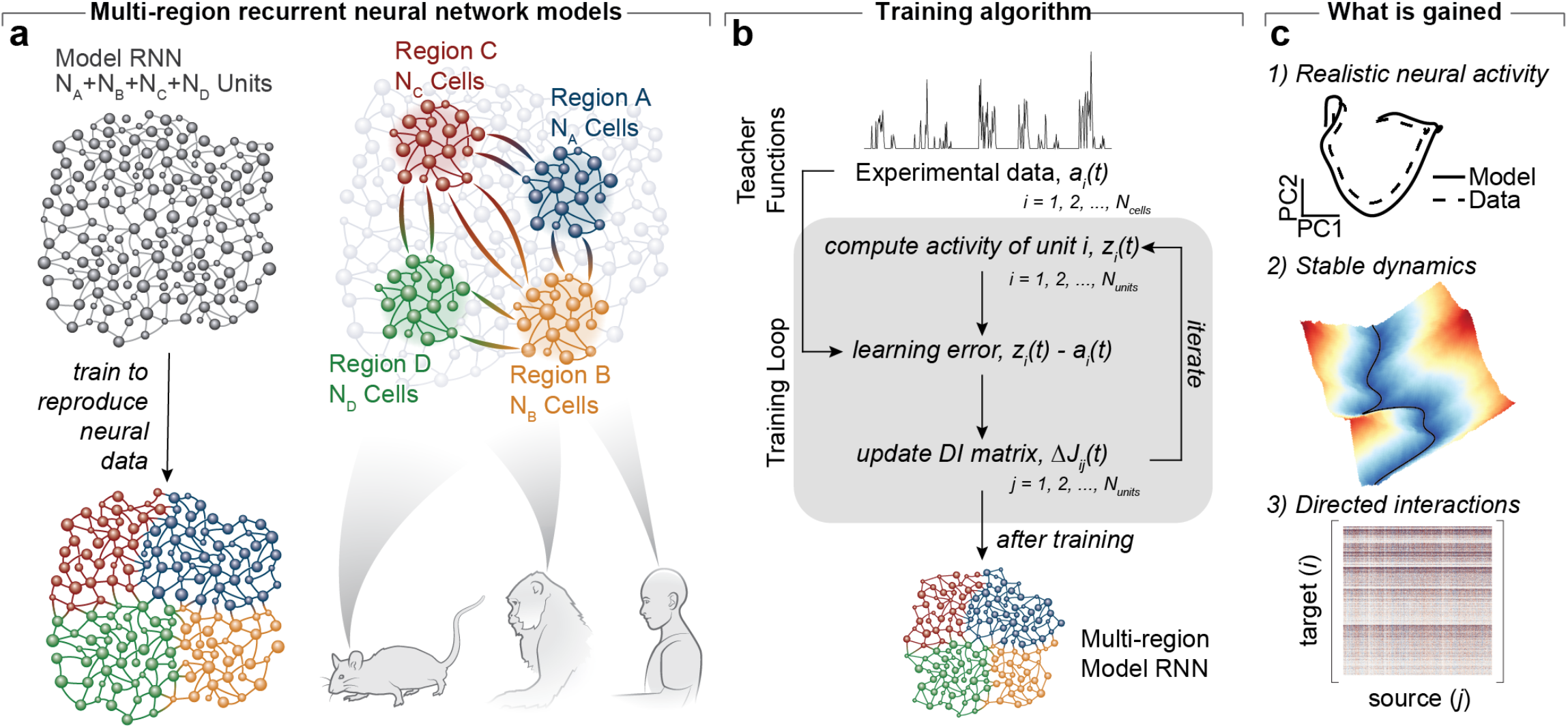
Data-constrained multi-region RNN design, training procedure, and outcomes. **(a)** CURBD is implemented through a Model RNN constrained from the outset by experimental data. Neural data from experiments in behaving animals (here, mice, monkeys, and human participants) are segmented into modules such as brain regions. A Model RNN is constructed such that each unit is trained to match a single experimentally recorded neuron from the full dataset of neural population activity from multiple interacting regions. **(b)** Training occurs where the connectivity matrix **J** of the Model RNN is modified over time until the activity of the RNN units match the experimental data. **(c)** This approach has several advantageous outcomes. 1) The model, after training, exhibits realistic neural dynamics consistent with experimental data. 2) The trained multi-region RNN produces a stable dynamical system. 3) The directed interaction matrix inferred by the trained Model RNN gives unique insight into the functional connectivity responsible for the observed dynamics in the data, including strength and type (e.g., excitatory or inhibitory and unidirectional or feedback projections between regions) of interactions, both within and across regions.

Training proceeds iteratively^10,17,19^ (**Figure 2b**; see Methods) in which the instantaneous linear error between each Model RNN unit (*z*_*i*_*(t)*) and the activity of its corresponding experimentally recorded neuron (*a*_*i*_*(t)*) is minimized. At each training step, the directed interaction matrix **J** is updated by Δ**J**, a function of this error (Equation 2; see Methods). Note that the Model RNN can be trained either from trial-averaged data aligned on relevant events or, when large numbers of simultaneously-recorded neurons are available, using single-trial or even continuous time-series data. Thus,

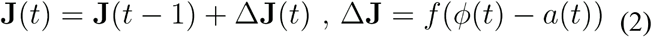

At this stage, to train the Model RNN, we do not need to incorporate assumptions about the identity of the modeled neurons, such as brain region (or cell type, cortical layer, etc). The Model RNN instead learns a single dynamical system that autonomously reproduces the entire sequence of multi-region experimental data using just an initial condition. In essence, after training we obtain an *in silico* model of the recorded brain regions that recapitulates the experimentally recorded multi-region data, but with crucial advantages (**Figure 2c**): i) the Model RNN natively generates realistic patterns of neural activity; ii) training tames the chaotic dynamics of the randomly initialized network^10^, ensuring that the trained network is dynamically stable; and iii) the model contains the directed interaction matrix that CURBD leverages to infer the currents between recurrently connected units both within and between regions. Since the Model RNN directly reproduces time-series neural data, this directed interaction matrix is an estimate of the functional interactions between each recorded neuron. These functional interactions are distinct from anatomical connectivity since they can include long-range and indirect effects such as neuromodulator release. Consequently, the currents in the RNN represent a functional estimate of the information exchanged between neurons in the recorded dataset, rather than a direct measure of physiological currents such as postsynaptic potentials.

The current into any one target unit *i=1*, 2, …, *N* can therefore be viewed as the sum of the activity of the *N* source units scaled by the respective interaction weights between the source units and the target unit (Equation 3).

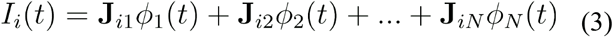

Since this is a dot product, all of the constituent source currents sum to reconstruct the full activity of units in the target region. In this paper, we focus on currents exchanged between brain regions. By restricting the summation in Equation 3 to source units from a specific region, we can isolate the currents into the target region from a specific source region (**Figure 1d**). In practice, based on labels applied to each experimentally recorded neuron, the matrix **J** can be broken into *M*^*2*^ submatrices, where *M* is the number of regions identified in the dataset, corresponding to all pairs of source/target interactions in the region (**Figure 1c**). Note that in this paper we assume that the region identities for each neuron are known *a priori* through anatomical labeling or other forms of clustering. This separation of currents can be considered as a decomposition of the activity of the target-region neurons based on the relative contributions of each source region. These source currents can be powerful tools to analyze existing neural data and help guide new experiments to dissect multi-region interactions. Direct analysis of the characteristics (e.g., strength, type, or timing) of the disparate current inputs can provide insight into the multi-region interactions that produce cohesive behavior. Additionally, the source currents inferred by CURBD provide a unique view into the inputs shaping neural population activity that are not easily observed experimentally.

## RESULTS

### Validation of CURBD on ground truth datasets

Since CURBD was designed to infer unobservable interactions in experimental datasets, we first validated the method in simulations where the ground truth inter-region currents are known. We created a generator model comprising three chaotic RNNs representing distinct regions with sparse inter-region connectivity (**Figure 3a**; see Methods). Region B was externally driven by a sequentially active population, Region C was externally driven by a population generating fixed points, and Region A was driven only through interactions with Regions B and C. We designed the simulation such that Region A was highly chaotic without clear representations of either external input (**Figure 3d**).

**Figure 3.**
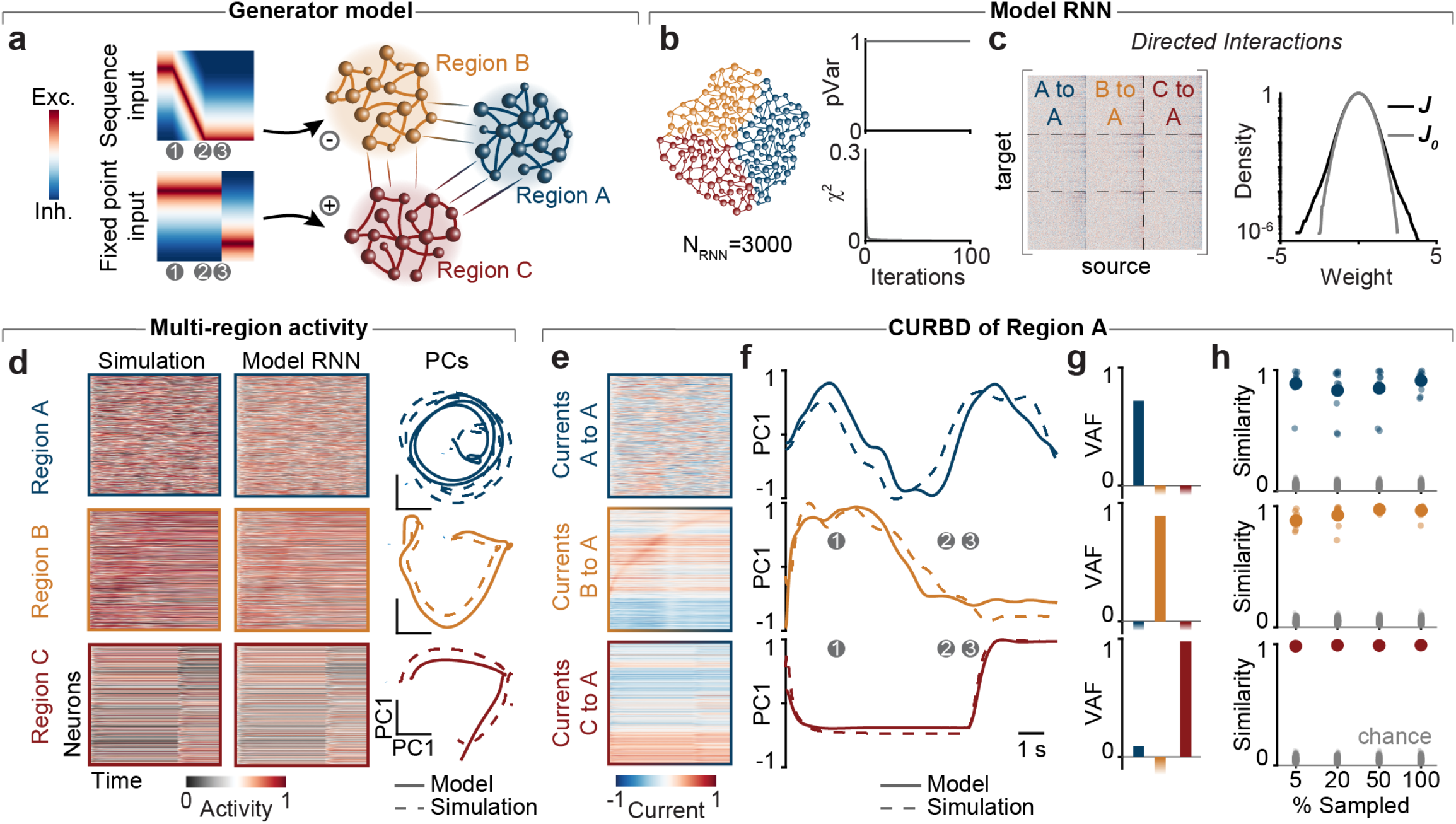
CURBD accurately isolates current sources in idealized, ground truth data from three interacting ‘regions’. **(a)** We generated three RNNs representing three distinct brain regions. Region B was externally driven by a sequentially active population, Region C was externally driven by a population generating fixed points, and Region A was driven only through interactions with Regions B and C. Time points 1-3 denoted by gray circles represent key time points in these external inputs. **(b)** We fit the Model RNN to the time-series data of all three regions comprising the generator model, reaching high variance explained (pVar=0.99) and low training error (*χ*^2^=7.4×10^−6^). **(c)** (Left) Training resulted in the directed interaction matrix describing the interactions within and between regions. (Right) Normalized distribution (log scale) of all weights in the Model RNN directed interaction matrix (**J**, black) compared with the randomly initialized matrix (**J**_**0**_, gray) **(d)** After training, the Model RNN accurately reproduced the single-unit activity of all three simulated regions (left), as well as the population trajectories in the leading principal components (PCs; right; solid lines: model; dashed lines: simulation). **(e)** The individual current sources into Region A showed qualitatively similar activation patterns to those in the source regions, even though these patterns were not apparent in the population activity of Region A. **(f)** The dominant PC of each source current (solid lines) and the ground truth inferred through the known connection weights in the generator model (dashed lines). Gray circles are the same key time points as in Panel a. **(g)** Variance Accounted For (VAF) for the first PC of each source current compared to the three ground truth source currents (VAF_AtoA_=0.72; VAF_BtoA_=0.89; VAF_CtoA_=0.98). **(h)** We repeated the simulation while randomly sampling different subpopulations of the 1000 unit networks ranging from 5 to 100% of the total units. Note that the 100% mark represents repeated training runs of the same neurons with different random initializations of **J**_**0**_. We quantified CURBD performance using a “similarity metric” of the current dynamics (see Methods). Small dots show the similarity from 10 different repetitions; large dots represent the average. Gray dots show the lower-bound obtained from 100 random shuffles of the ground truth dynamics.

We trained a single Model RNN (**Figure 3b**) to match the simulated data from the generator model (**Figure 3c-d**; see Methods). We hypothesized that CURBD would accurately infer the inputs to Region A from source Regions B and C despite the chaotic nature of the population activity observed in Region A. Using the submatrices of the directed interaction matrix (**Figure 3b**), we decomposed the activity of Region A into the currents from each source region. These currents showed qualitatively similar activation patterns to those in the source regions (**Figure 3e**), even though these patterns were not apparent in the population activity of Region A. Since the true connectivity of the simulated network was known, we also computed the ground truth currents into Region A. We summarized the population-wide currents from each source using principal components analysis (PCA) and compared the CURBD output to ground truth using Variance Accounted For (VAF; see Methods). CURBD accurately reconstructed each current source driving Region A (**Figure 3e**). We then compared the performance of CURBD to canonical correlation analysis (CCA), which can identify individual subspaces that capture the shared dynamics between pairs of regions. We found that CCA did not infer the ground truth currents (**Figure S2**) due to the recurrence in the network^27^.

We adapted the simulation to test the practical limits of CURBD. In real datasets, experimenters typically only have access to a small percentage of neurons in a given region. We repeated the simulation to test whether CURBD is effective when the brain regions are partially sampled by training the Model RNN to target only a subset of units. We computed a “similarity metric” (see Methods) that could compensate for the different number of recorded neurons^12,28,29^ to compare the current inferred by CURBD and the ground truth. CURBD accurately estimated the current dynamics even when the network was highly undersampled, as low as 5% of the population (**Figure 3h**). We then designed a second simulation to explore the regimes where CURBD succeeds. We simulated two recurrently connected RNNs, each receiving sinusoidal inputs of different frequencies (**Figure S3**). Since the sinusoidal inputs can mix with the ongoing chaotic dynamics in recurrent networks, they provide a more challenging paradigm to assess CURBD. We found that CURBD was most effective when the intrinsic dynamics of the two RNNs were distinct, with sparse inter-region connectivity (**Figure S3g**). These simulated ground truth datasets illustrate that CURBD can accurately infer unobserved source currents between multiple brain regions under a variety of conditions. In the following sections, we apply CURBD to multi-region experimental recordings to infer brain-wide currents in behaving animals.

### CURBD untangles brain-wide currents during spontaneous movement in mice

Optical recording of fluorescence from genetically encoded calcium sensors allows experimenters to simultaneously track the activity of thousands of neurons from across the brains of behaving animals. Here, we demonstrate that CURBD untangles behaviorally relevant source currents from large-scale, multi-region calcium imaging datasets. Mice expressing GCaMP6s^30^ were allowed to run spontaneously on top of an air-supported ball in complete darkness (**Figure 4a**). Using a large field-of-view two-photon microscope^31^, we imaged neural activity simultaneously from four regions (**Figure 4a-b**): primary visual cortex (V1), secondary motor cortex (M2), posterior parietal cortex (PPC), and retrosplenial cortex (RSC). Together, these regions contribute to a brain-wide circuit governing navigation, decision-making, and movement^32–34^. Mice exhibited spontaneous bouts of running behavior, measured as rotations of the air-supported ball (**Figure 4c**), with complex patterns of neural activity across all four brain regions during these bouts. Consistent with recent studies^5,6^, we observed a high degree of activity even in V1 despite the fact that the mice received no visual input, highlighting the distributed nature of behavior-related activity throughout the brain.

**Figure 4.**
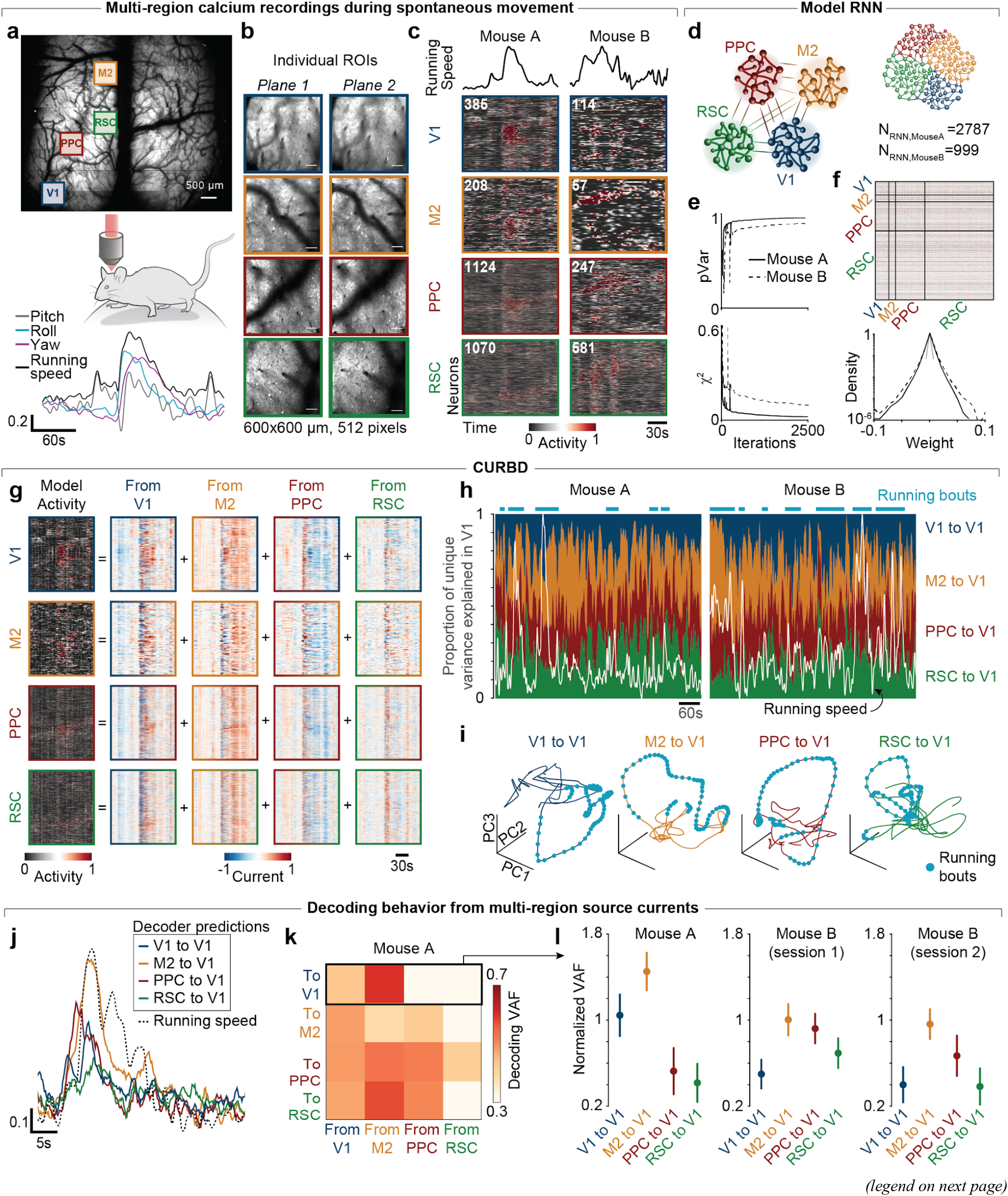
Isolating source currents from multi-region calcium recordings in mice. **(a)** We recorded neural activity from four brain regions in two mice expressing GCaMP6s. Mice were head-fixed on an air-supported ball in complete darkness. Running was tracked using sensors recording the pitch (gray), roll (cyan), and yaw (magenta) velocities of the ball; the total magnitude of the three signals, combined, is summarized as running speed (black). **(b)** We imaged two planes from each brain region (regions of interest, ROIs). **(c)** Behavior and neural population activity from the four regions during a brief period of spontaneous running for two mice. Text inset denotes the number of recorded neurons in each region. **(d)** We used ten consecutive minutes of recordings to fit a Model RNN for both mice. **(e)** Proportion of variance explained (pVar) and training error (*χ*^2^) for the RNNs trained to match data from Mouse A (solid lines) and two sessions from Mouse B (dashed lines). **(f)** (Top) Example directed interaction matrix for Mouse A. (Bottom) Normalized distribution of interaction weights (log scale) for the three Model RNNs before (gray) and after (black) training. **(g)** CURBD decomposition for Mouse A. (Left) Heatmaps of RNN unit activity for the four regions. (Right) Heatmap of current decomposition for each of the sixteen source currents capturing all possible inter-region interactions. **(h)** The proportion of unique variance explained by each source current of the total V1 activity. Running speed is overlaid in white, and cyan lines indicate running bouts. **(i)** V1 source current trajectories in the three leading PCs for Mouse A. Cyan dots denote time points at which the mouse running speed was above a threshold ball speed. **(j)** We used linear decoders to predict running speed from each source current. Example decoding predictions (colored lines) of the measured running speed (dashed line) for the four source currents into V1 for Mouse A. **(k)** VAF for all sixteen source current decoders for Mouse A. **(l)** Decoder performance (mean and standard deviation across 1000 random cross-validated test sets; see Methods) for source currents into V1 for Mouse A and two sessions from Mouse B.

We hypothesized that CURBD could isolate the sources of behavioral information in regions such as V1. We trained Model RNNs to reproduce the neural data from the four recorded regions (**Figure 4d-f**). Applying CURBD, we identified strikingly different patterns of excitation and inhibition during running bouts for the sixteen source currents (**Figures 4g, S4a,c**). Our analysis focused on the currents into V1 seeking to identify sources of signals related to running. We computed the relative variance explained by each source current of the full V1 population activity (**Figure 4h**; see Methods). We predicted that currents from M2 and PPC, which are closely involved in planning and producing behavior^32,35^, would increase during bouts of running. However, we saw no clear relationship between the variance captured by each source current and the running speed. Instead, source currents between the four brain regions occurred in similar proportion.

To analyze the population-wide dynamics of these currents, we computed low-dimensional neural manifolds^7,36^ spanning each source current using PCA. The trajectories within these manifolds (**Figures 4i, S4b,d**) capture the dominant dynamics of each source current into the target region. Studying the dynamics of the source currents to V1 (**Figure 4i**), we observed that while M2 and PPC currents showed large deviations in their trajectories, RSC to V1 currents did not greatly change during running bouts. These observations suggest that information related to ongoing behavior may arrive in V1 selectively from M2 or PPC. To test this quantitatively, we built linear decoders predicting running speed based on each source current to V1 (see Methods). Comparing decoders trained using the currents into V1, we found that the currents from M2 contained the most information about running speed (**Figures 4j-l, S5**). These results illustrate the potential of CURBD to untangle the complex, multi-regional interactions underlying behavior using RNNs based on multi-region calcium imaging data.

### CURBD separates inter-region interactions from spiking data collected during Pavlovian conditioning in monkeys

We next applied CURBD to population spiking activity acquired by electrophysiological recordings in macaques. Many multi-region population datasets obtained by electrophysiology are constructed using “pseudopopulations” where neurons recorded at different times are pooled together by averaging across repetitions of the same condition. We thus aimed to demonstrate that CURBD could infer currents from pseudopopulation datasets. We obtained neural data from two monkeys (*Macaca mulatta*) performing a Pavlovian conditioning task (**Figure 5a**). The monkeys learned to associate three conditioning stimuli with three different reward levels of increasing desirability: no reward (CS-), water, and juice. After a brief anticipatory delay, the monkeys received the expected reward. Using intracranial electrodes, we recorded from neurons in the amygdala, subcallosal anterior cingulate cortex (ACC), and rostromedial striatum. These regions are known to be important for reward processing and affective behaviors^37^. Since we could record only small numbers of neurons on a given session, we constructed pseudopopulations of 343 neurons for Monkey D and 199 neurons for Monkey H by averaging neural activity across all trials for each condition on each session. All three regions displayed condition-specific sequence-like activity^38^ (**Figures 5b, S6d,i**).

**Figure 5.**
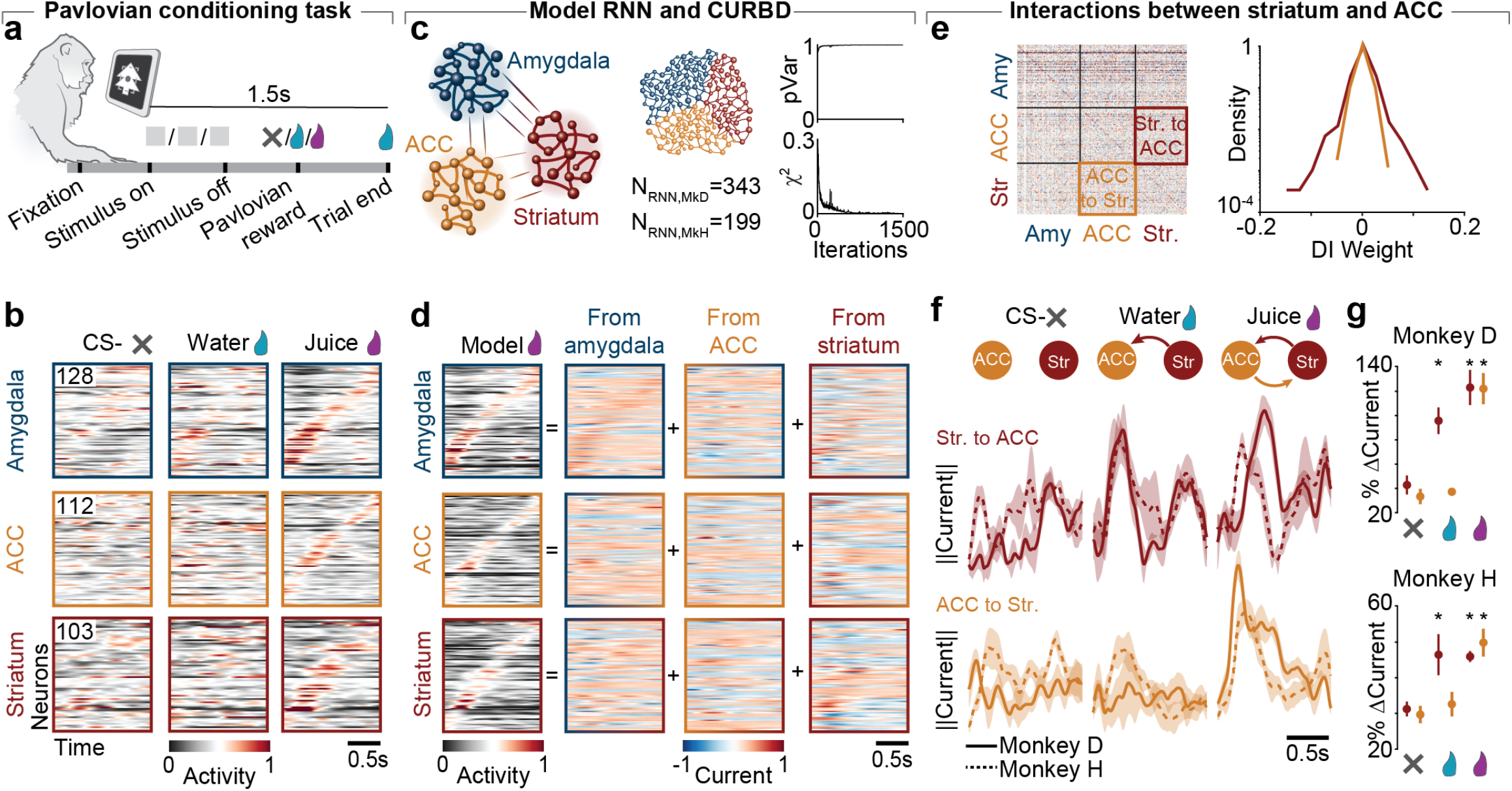
Current-based decomposition of three-region pseudopopulation recordings in macaques. **(a)** Macaque monkeys performed a Pavlovian conditioning task where one of three stimuli associated with no reward (unconditioned stimulus), water, or juice were presented for 1 second. The associated reward was delivered after a short delay (0.4-0.6 seconds) then a second water reward signified the trial end. **(b)** Trial-averaged firing rates in the pseudopopulation dataset for Monkey D for the amygdala, subcallosal ACC, and striatum during the unconditioned stimulus (left, inset number denotes neuron count in each region), water stimulus (middle), and juice stimulus (right). Neurons in each region are aligned on the presentation time of the stimulus and sorted according to their time of peak activity in the juice condition. **(c)** (Left) Schematic of Model RNN. (Right) Proportion of variance in the neural population explained by the model (top, pVar) and training error (*χ*^2^, bottom) as a function of the number of training iterations. **(d)** CURBD of activity in each region for the juice trials. Left heatmaps show the full Model RNN activity. Remaining heatmaps show the decomposition for each of the sixteen source currents capturing all possible inter-region interactions **(e)** (Left) Directed interaction matrix for an example Model RNN from Monkey D. (Right) Distribution of weights (log scale) in the striatum to ACC (red) and ACC to striatum (yellow) submatrices. **(f)** Magnitude of bidirectional currents from striatum to ACC (red, top) and ACC to striatum (yellow, bottom) during presentation of the three stimuli. Solid line: Monkey D; dashed line: Monkey H. Error bars: standard deviation across five different random initializations of the Model RNNs. Schematics (top row) summarize the dominant source currents inferred by CURBD—magnitude and directionality—between the two regions. **(g)** Statistical summary of the percent change in total current in the first 1s of each condition compared to the mean in the unrewarded condition. Points represent mean and lines s.e.m. *: significance at p<0.05, t-test.

We trained Model RNNs to reproduce the neural data for each monkey (**Figures 5c, S6a,b,f,g**). The RNNs accurately learned the neural dynamics of the three regions even though the neurons were not simultaneously recorded. Inspecting the nine source currents, we saw that CURBD uncovered distinct dynamics for each region in the circuit (**Figure 5d**). One notable advantage of CURBD is that it can infer directed inter-region currents to determine, for example, whether the interactions between two regions are reciprocal or feedforward. Since ACC directly projects to rostromedial striatum^39^, we focused our analysis on these two regions.

Intriguingly, we found that the strength of interactions between striatum and ACC were asymmetric (**Figure 5e, Figure S6c,h**). Since the inferred currents are the product of both the interaction strength and the source region activity, we further studied the asymmetries in the bidirectional interactions using the total magnitude of current between the two regions (**Figure 5f-g**). We saw strong currents from Striatum to ACC following the water stimulus, but no corresponding current from ACC to Striatum. However, on the juice trials, we saw strong bidirectional currents.

Crucially, the currents inferred through CURBD were consistent across both monkeys, as well as across five different random initializations of the Model RNN.

Since the pseudopopulations are constructed *post hoc*, their size and the specific neurons that are chosen for inclusion in the population can be arbitrary. We tested whether CURBD infers the same population-wide current dynamics with pseudopopulations constructed by sampling different subsets of the total population of neurons. We randomly subsampled the available neurons from each region to create pseudopopulations of different sizes (between 60% and 90% of the total) and performed CURBD. We computed the similarity metric as above for each of the nine source currents, comparing the inferred currents at each sampling percentage to the currents inferred when using the full population. We found a high degree of similarity in the identified currents even when using just 60% of the available neurons (**Figure S7**). Thus, CURBD can be readily applied to pseudopopulations comprising non-simultaneous recordings, yielding robust estimates of the interactions between regions.

### CURBD applied to single-cell spiking data from humans during memory retrieval

We next applied the method to cellular resolution, multi-region, spiking electrophysiology recordings from humans. Five participants performed a set of two memory tasks in eight interleaved blocks (**Figure 6a-c**, see Methods)^40^. In the first, participants categorized images based on high level sensory features. In the second, participants were presented with an image and reported whether or not they had seen the image before. As the participants performed this task, we recorded the activity of neurons in two frontal cortical regions—pre-supplementary motor area (preSMA) and dorsal anterior cingulate cortex (dACC)—and the hippocampus and amygdala (H/A) using hybrid depth electrodes^41^. Using the same procedure as in the monkey dataset, we constructed pseudopopulations from neurons recorded from between two and five sessions in each participant. Since some participants had few recorded neurons from either hippocampus or amygdala, we combined them for later analyses^40^. Memory retrieval is believed to be mediated by interactions between frontal cortices and the H/A^40^. Our goal was to demonstrate that CURBD could separate currents related to the memory retrieval and memory formation within these regions. Thus, we focused our analysis on the memory task, where participants accessed their memory after viewing each image and instantiated a new memory following a novel image.

**Figure 6.**
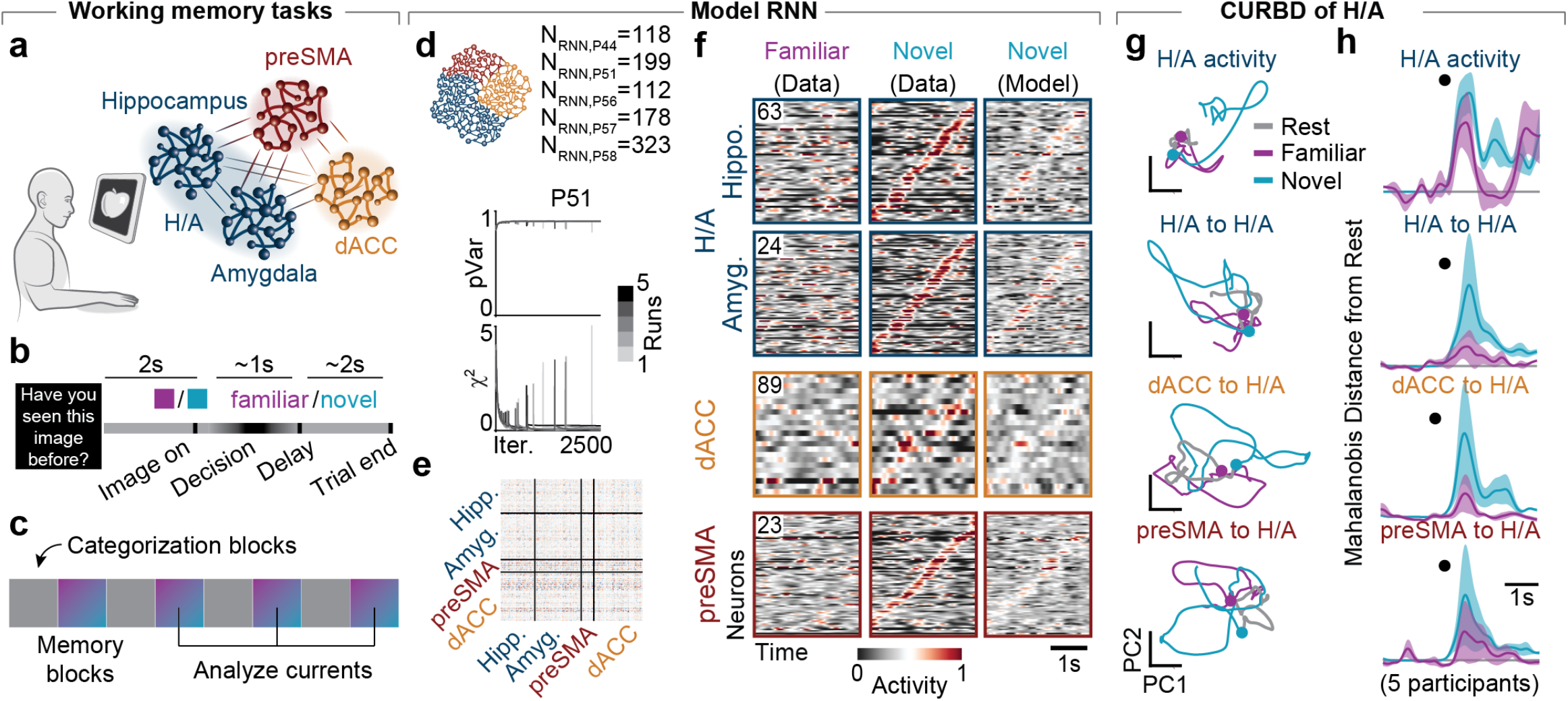
Inferring source currents between four regions in humans performing a memory-retrieval task. **(a)** Human participants were implanted with depth electrodes to record single-unit spiking activity from neurons in the hippocampus and amygdala (combined and abbreviated H/A), pre-supplementary motor area (preSMA), and dorsal anterior cingulate cortex (dACC). **(b)** Trial structure during each memory block. After a two-second baseline period, a familiar or novel image was presented. Participants reported whether they had previously seen the image (familiar) or not (novel). **(c)** Each experimental session comprised eight blocks. In odd blocks the participants categorized images; in even blocks, as schematized in Panel b, the participants reported whether a presented image was novel or familiar. We used data from blocks 4, 6, and 8 to compare familiar and novel stimuli when task performance was highest. **(d)** Training performance (pVar and *χ*^2^) for Model RNNs in P51, shades of gray denote five different random initializations (runs). **(e)** Directed interaction matrix of a Model RNN trained to match data from P51. **(f)** Pseudopopulation activity during the memory task (Block 4) from P51 following familiar (left, inset number denotes neuron count) and novel (middle) images, and the corresponding Model RNN activity on novel trials (right) for the four regions. Neurons within each region are sorted based on the time of peak activity in the recorded data on the novel trials. **(g)** Population trajectories projected onto the leading two PCs for H/A neurons during the pre-stimulus baseline period (Rest, gray) and in response to familiar (magenta) and novel (cyan) stimuli, and the source current trajectories within H/A for the two types of stimuli. Dot indicates the state at the time of stimulus onset. **(h)** Mahalanobis distance from the pre-stimulus rest period computed over time for each source current into H/A. Dot indicates time of stimulus onset.

We fit Model RNNs to the pseudopopulation datasets from each participant (**Figures 6d-f, S8**) to estimate the directed interaction matrices. We then performed CURBD to infer the currents driving H/A following presentation of familiar or novel images. Inspecting trajectories in the first two PCs of the full H/A activity and each source current, we saw distinct current patterns between the categorization and memory tasks after image presentation (**Figure S9**). Within the memory task, the currents within the circuit also changed between novel versus familiar image conditions, with familiar images causing a small response in the frontal cortex to H/A currents and novel images causing a large response in all currents. We quantified these effects using the Mahalanobis distance from the cluster of resting state activity (**Figure 6h**; see Methods).

Across all five participants, CURBD identified a substantial change in currents (relative to baseline) from preSMA to H/A when viewing familiar images, and smaller changes in the other source currents. Viewing novel images, on the other hand, caused large sustained currents throughout the whole network. These results suggest a specificity in the inter-region interactions inferred by CURBD: frontal cortex provides input to H/A during memory retrieval, while the remaining pathways are recruited following a novel image to encode new information into memory.

## DISCUSSION

### Advantages of CURBD

Typical data analysis approaches study population activity from the perspective of the experimentally measured outputs from a neural circuit (e.g., action potentials through electrophysiology or calcium fluorescence signals through imaging). Using dimensionality reduction techniques^7^, we can estimate low dimensional neural manifolds^36,42^ embedded in the space of total population activity. Neural manifolds are defined by patterns of covariation between neurons in measured population activity. However, the covariance observed in neural populations is shaped by the inputs driving that population^43^. CURBD offers a unique view of neural activity by decomposing experimentally measured population activity into such inferred inputs or ‘source currents’. Rather than identifying a single manifold capturing the measured outputs of active units within a given region, we can use the source currents to compute a separate manifold, one for each source current inferred. Therefore, CURBD allows us to reconceptualize population activity as numerous manifolds embedded in the space of neural activity, each capturing the dynamics of a single, isolated source of input.

CURBD addresses several gaps in commonly applied computational approaches for analyzing experimental data enabled by new technologies for monitoring large scale neural activity from multiple interacting brain regions ^31,44–48^. Common methods to study interactions between brain regions such as linear regression^23,26^, CCA^49^, constrained dimensionality reduction^24,25^, generalized linear models (GLMs)^23^, or Granger causality^50^ rely on correlative analysis of neural data, posing several challenges. First, correlation-based inference of functional connectivity cannot distinguish between correlations that arise from common inputs and those that arise from other types of interactions between regions, though these can be partially accounted for by incorporating additional covariates^23,51^. Second, correlation alone does not provide directionality, though careful assessment of spike latencies can provide some insight into possible directional effects^52^. Third, correlative analyses typically describe interactions between two regions and are difficult to extend to data from multiple interacting regions, though recent work on switching dynamical systems shows promise^53,54^. CURBD addresses these limitations by building and analyzing RNNs that are trained to match the entire time-series from experimentally collected data^2,86^. Thus, CURBD explicitly models the recurrence between all recorded neurons, capturing all possible multi-region interactions in the dataset. This allows us to, in an unbiased way, capture the directionality and magnitude of the interactions within and across regions that are responsible for the observed neural dynamics. Furthermore, the directed interaction matrix inferred from the trained multi-region RNN is asymmetric, allowing directional estimates of the inferred functional interactions (e.g., **Figure 5e**). Lastly, since CURBD concurrently models all multi-region interactions, it scales natively to arbitrarily large datasets with any number of regions, even to whole-brain recordings available from *Caenorhabditis elegans*^*55*^ and larval zebrafish^19,44,45^. In contrast to dynamic causal modeling^56^, CURBD does not necessarily require known perturbations or inputs, and can flexibly model any dataset. CURBD also natively models the inherent dynamical stability of the neural data. This biologically-relevant constraint leads to more specific and meaningful solutions.

### Interpreting directed interactions inferred from RNNs constrained directly by data

While CURBD estimates multi-region interactions by incorporating recurrence within and between regions and dynamical stability, these interactions should not be considered causal relationships. Additionally, the directed interactions estimated by the Model RNN need not relate to actual synaptic connectivity. While a direct monosynaptic connection between two neurons should contribute to a strong directed interaction weight, strong interactions could arise indirectly as well^57^. Polysynaptic pathways (including those involving neurons or brain regions that were not experimentally observed) or triggering neuromodulator release could enable one neuron to exert an influence on other neurons that would manifest as an inferred directed interaction weight^19,58^. Additionally, brain-wide state changes that impact distributed neural circuits—such as those induced by stress^19^, depression^59^, or even glia^60^—could lead to strong functional relationships between recorded neurons.

### Model RNNs underlying CURBD are a dynamical system

The Model RNNs used for CURBD are specific, learned dynamical systems that capture the essential features of theneural dynamics from the data they were trained to match based on an initial condition. This facet represents a difference between CURBD and other approaches that seek generative models of the neural dynamics^21^. However, even though our models here are typically fit only to single instantiations of data^61^, we identify consistent solutions from one iteration to another, for instance, at the level of statistical distributions of groups of interaction weights (e.g., **Figure S6**) as well as at the level of currents inferred by CURBD (e.g., **Figure 5f**). Furthermore, since the inter-region currents inferred by CURBD rely on the product of the directed interaction weight matrix and the activity, the estimation noise in different realizations of the matrix are averaged out. Therefore, the currents identified by CURBD are robust to different random initializations of the directed interaction matrix, allowing for consistent solutions under a variety of initialization conditions, as well as to different random subsamples of the modeled neurons.

Ultimately, the Model RNNs underlying CURBD should be considered as a model of the data itself—an *in silico* representation of the experiment. This model enables a deeper dive into the experimentally measured data using the directed interaction matrix or currents due to inter-region interactions which we cannot access experimentally^62^. Our current approach assumes that a single directed interaction matrix captures the dynamics for the whole duration of the data. Factors such as learning^63,64^ or behavioral state changes^19^ could change the dynamical rules governing the interactions among different neural populations in vivo. If such state changes are identified, they can be addressed by fitting different Model RNNs on different samples of data (e.g. periods of time, task conditions). The final currents can then be fully reconstructed by essentially “stitching together” the currents inferred by Model RNNs fit to each set of samples. More elegantly, the training process could also be modified to identify state changes and adjust the directed interaction matrix over time in a partially unsupervised, adaptive manner.

### Additional uses and extensions of CURBD

The multi-region Model RNNs employed in the applications above made no assumptions about the structure of the directed interaction matrix or inter-region connectivity. Instead, we allowed the neural networks to opportunistically, through the process of training, construct solutions that recapitulated the essential dynamical features in the multi-region experimental data. In biological systems, there are numerous anatomical constraints that could be incorporated into the model in the future. For example, the effect of a given neuron on its numerous downstream targets is typically either excitatory or inhibitory^65^. This constraint could be incorporated into the learning rule such that columns of the directed interaction matrix are restricted to have either all positive or all negative weights. Additionally, while we allowed our RNNs to be weighted all-to-all in the inter-regional interactions (the off diagonal submatrices), inter-regional connections in biological brains are highly structured. For example, long-range connections between regions are likely more sparse than within a local population^66^. Such sparsity could be induced in an unsupervised manner by applying an L1 norm on the weights of specific submatrices in the cost function. Brain-wide connectomics data^57,67,68^ could also be leveraged to build a prior into the directed interaction matrix about which pathways should be directly connected. Lastly, we trained the Model RNNs using rates estimated from the neural recordings. Future extensions of CURBD could allow more temporally-precise directed interaction estimates by incorporating spiking statistics models into the training.

In the present work, the region identity of each experimental neuron was known using anatomical landmarks or electrode implantation site. This knowledge allowed us to readily divide the directed interaction matrix into region-specific blocks. However, we predict that in future work Furthermore, we predict that CURBD can be extended to provide a basis for functional clustering that goes beyond anatomical designations by applying clustering or tensor decomposition methods^69^ directly to the currents inferred by the Model RNN into each target unit. CURBD could be used in an unsupervised manner to find relevant population designations based on functional distinctions and their interactions with other neurons, identifying functional submodules within single regions^70^ and identify brain-wide functional circuits.

We used calcium fluorescence and spiking activity from single, identifiable neurons to constrain the Model RNNs for CURBD, but the possible use cases of the general approach are not confined to cellular resolution data. Model RNNs can be fit to non-cellular resolution data, such as multi-electrode local field potential recordings. Furthermore, other types of relevant experimental data or conditions can be incorporated as additional constraints on the Model RNNs during training. Behavioral data such as body posture derived from modern pose detection methods^71,72^ could be incorporated into the training process to help account for unobserved common inputs related to that behavior^73^. Static labels representing experimental metadata (behavioral task, stimulus condition, etc.) could also be incorporated to help compensate for brain-wide state changes. These measurable external signals could be targeted to all recorded neurons, or a specific subset (e.g. brain region, cell type) if such constraints are known. Importantly, all of the extensions described above do not change the fundamental principles underlying CURBD, which at its core, relies on straightforward matrix multiplication. They only serve to provide a more constrained estimate of the biological system’s directed interactions.

The power of CURBD lies in harnessing the ability to flexibly engineer multi-region RNNs based on a broad range of time-series data from various experiments, as we have exemplified with the four applications presented here. There is often a remarkable conservation of structure and function throughout evolution and across species producing a certain behavior even with divergent phylogenetic trees^74^. Therefore, understanding the commonalities (or unique differences) in identified mechanisms across different species will be critical to uncover fundamental principles of neural computation^1,75^. This requires an analytical framework such as CURBD that robustly and flexibly scales across a range of different experimental—e.g., methodologies, levels of granularity, sampling densities, and spatiotemporal resolutions—such as those encountered when comparing different species ranging from smaller, highly sampled nervous systems (e.g., *Caenorhabditis elegans, Drosophila*, larval zebrafish) to larger, less sampled brains (e.g., rodents, non-human primates, and humans). Thus, CURBD provides a powerful new approach for comparative studies over time, across individuals, scales of neural function, or even species.

## ACKNOWLEDGEMENTS

C.A. is supported by a Louis Perry Jones, an Alice and Joseph Brooks, and a Mahoney Postdoctoral Fellowship. S.S. is supported by an European Molecular Biology Organization (EMBO) Postdoctoral Fellowship. U.R. is supported by the National Institutes of Health (NIH) (R01 MH110831 and U01 NS117839) and National Science Foundation (NSF) (BCS-1554105). P.R. is supported by a National Institute of Mental Health BRAINS award (R01MH110822, R01MH118638) and a young investigator grant from the Brain and Behavior Foundation (NARSAD). C.D.H. is supported by an NIH Director’s Pioneer Award (DP1 MH125776) and NINDS R01 NS089521. K.R. is supported by NIH BRAIN Initiative (R01 EB028166), James S. McDonnell Foundation’s Understanding Human Cognition Scholar Award, and NSF FOUNDATIONS award (NSF1926800). We would like to thank Dr. Juan A. Gallego, Dr. Raeed H. Chowdhury, Dr. Larry F. Abbott, Dr. Sheila Cherry, and Dr. Christian D. Marton for comments on earlier drafts of this manuscript. We are indebted to and inspired by Dr. C. R. Rajan and Prabha Rajan for sharing their undying love of learning.

## AUTHOR CONTRIBUTIONS

M.G.P. and K.R. conceived of the method. M.G.P. analyzed datasets and generated figures. M.G.P. and K.R. wrote the manuscript. M.G.P., C.A., S.S., C.M., J.M., U.R., P.R., C.D.H., and K.R. edited the manuscript. E.C. ran simulations and implemented code. C.A., S.S., and C.D.H. collected the mouse dataset. M.E.Y., C.M., and P.R. provided the monkey dataset. J.M. and U.R. provided the human dataset.

## METHODS

### Code availability

All modeling and analysis in this manuscript was done in Matlab (The Mathworks, Inc.). Matlab and Python code to train multi-region Model RNNs based on multi-region experimental recordings and perform CURBD using the inferred interactions is available at: https://github.com/rajanlab/CURBD.

### Multi-region recurrent neural networks

#### Network elements

Network models represent real biological circuits, but they do so with different levels of fidelity. We constructed Model RNNs that are directly constrained by experimentally-obtained time-series neural data. Each network unit or model neuron indexed by *i* is described by a total current *x*_*i*_, and an activation function, *ϕ*(*x*_*i*_), a nonlinear function of *x*_*i*_, where *i=1*, 2, …, *N* is the total number of units in the network. Each variable *x*_*i*_ obeys the following equation:

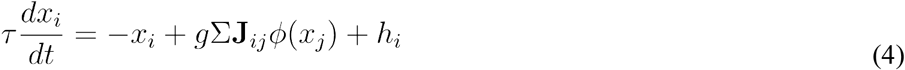

where *h*_*i*_ is its external input to the unit, **J** is a heterogeneous matrix of recurrent connections, and *τ* is the unit’s time constant selected to match the expected temporal dynamics of the data to be modeled (for the datasets in this manuscript, see **Table 2** below),. The control parameter *g* determines the strength of the recurrent connections, and thus whether (*g*>1) or not (*g*<1) the network produces spontaneous activity with non-trivial dynamics^76–78^. We set *g*=1.5 here, though in practice we observe qualitatively similar results for a range of values provided *g* is sufficiently large to facilitate chaotic dynamics in the network. The network equations are integrated using the Euler method and an integration time step, *Δt*_*RNN*_. Note that the network integration can occur with a finer time step than the sampling rate of the experimental data to be modeled (*Δt*). This allows for smoother dynamics when the experimental data may be sparsely sampled. We use *ϕ*(*x*_*i*_)*=*tanh(*x*_*i*_), but other saturating nonlinearities, such as sigmoids, have been explored in related work (see Refs. ^17^ and ^77^). This ensures that the firing rates go from a minimum of -1, which we conceptualized as a background rate, to a maximum at 1. The function also retains a maximum gradient at *x*=0.

Recurrent weights carrying inputs onto a target unit *i=1*, 2, …, *N* from its source partner *j*, **J**_ij_, which are the elements of the matrix **J**, are either fixed or modifiable (plastic), depending on how much structure is introduced into the connectivity of the initially disordered network. We introduce no *a priori* structure in **J**, allowing all elements to be modified during training. **J**_ij_ can potentially be modified by a number of different learning rules; here (see below). A crucial advance from previous modeling studies involves using a block-diagonal **J** in which each block represents the recurrent connections within each brain region being considered, and the off-diagonal blocks, the inter-region projections to and from them. In this way a two-region Model RNN has two blocks on the main diagonal relating each region to itself, and two regions on the off-diagonal relating each region to the other (e.g., **Figure S3b**). Following this pattern, multi-region Model RNNs are initialized with **J** matrices containing more than 2 blocks (e.g., **Figures 5e**).

Typically, the initial, untrained directed interaction matrix **J**_0_ is constructed to be the same size as the number of neurons in the dataset to be modeled, though larger networks in which different subsets of weights are modified in a data-dependent manner have been explored previously in Ref. ^17^. Here, each Model RNN unit is matched to one recorded neuron from the respective experimental dataset. The individual weights in **J**_0_ are initially chosen independently and randomly from a Gaussian distribution with mean and variance given by <J_0_>=0 and *<J*_*ij*_*>*^*2*^_*J*_*=g*^*2*^*/N*. Ultimately, the elements of **J** will be modified by the training algorithm until the activity of the model RNN’s units autonomously produce neural data consistent with the experimental recordings.

#### Design of external inputs

In real brains, neural populations are constantly driven by external and inter-regional inputs that we cannot always observe. To mimic this effect, we modeled background inputs that are uncorrelated with the relevant behavior we are studying. The external inputs to the units in the Model RNN, denoted by *h(t)*, are generated from filtered and spatially delocalized white noise that is frozen, using the equation:

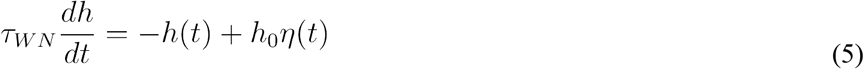

where *η* is a random variable drawn from a Gaussian distribution with 0 mean and unit variance, and the parameters *h*_*0*_ and *τ*_*WN*_ control the scale of these inputs and their correlation time, respectively. We use *h*_*0*_=1 and *τ*_*WN*_=0.1 in this paper. There are typically as many different inputs as there are model neurons in the network, with individual model neurons receiving the same input on every simulated trial.

#### Model RNN training

During training, the activity of individual units in the Model RNN, say the firing rate *ϕ*(*t*), are compared directly to teacher functions derived from the experimentally-recorded neurons, denoted by *a*_*i*_*(t)*. This gives an error function for each Model RNN unit:

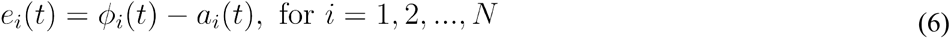

The activity of each *i*th target unit can also be computed as:

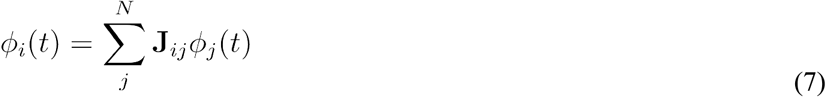

where *ϕ*_*j*_*(t)* is the firing rate of the *j*th source neuron (*j=1*, 2, …, *N*) connected to it through the recurrent weight **J**_ij_. During training, the elements in the directed interaction matrix **J** undergo modification at a rate proportional to three factors: i) the error term computed above; ii) the “presynaptic” or source firing rate of each neuron; and iii) a matrix **P** with *N*^*2*^ *pN x pN* elements. **P** is defined mathematically as the inverse cross-correlation matrix of the firing rates of units in the network, such that its elements **P**_ij_ are given by:

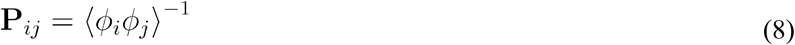

The matrix **P** keeps track of correlations in the firing rate fluctuations across the network at every time step, and is computed for all *i=1*, 2,…, *N* target units and *j=1*, 2, …, *N* source units.

Training proceeds iteratively as schematized in **Figure 2b**. At each time step, *t*, for *i=1, 2*,…, *N* target units, the corresponding elements of **J** are adjusted from their values at the previous time step (*t-1*) according to:

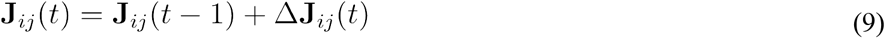

where the update term is computed according to Refs. ^10,79^:

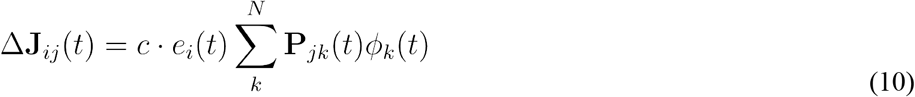

The scaling term *c* is computed according to:

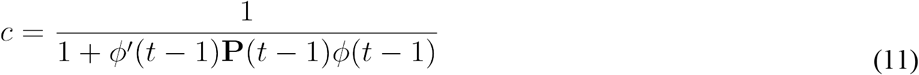

The error for each RNN unit *i* compared to its target, *e*_*i*_(*t*), is computed as:

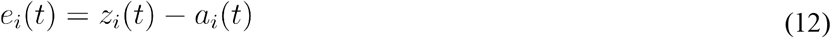

It is not generally necessary to calculate the matrix **P** explicitly. Instead, **P** can be updated iteratively according to Ref. ^79^:

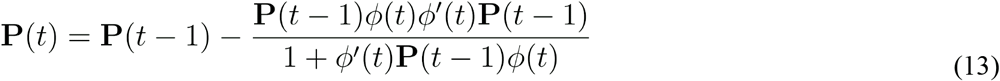

The matrix **P** is initialized to the identity matrix scaled by a factor *P*_*0*_ which controls the overall learning rate. In practice, training is most effective when *P*_*0*_ is set to be 1 to 10 times the overall amplitude of the external inputs (*h*_*0*_).

Since the learning algorithm updates **J** at each time step, high performance could be observed during training even when the algorithm has not fully converged. Thus, after training for a fixed number of iterations (typically between 1500 and 3000 iterations for model RNNs based on experimental neural datasets), we disabled the training for a few additional iterations to compute and evaluate the final goodness of fit. We assessed the quality of the fit and convergence using two metrics: 1) the training error (**χ**^2^) between the Model RNN rates and the teacher functions derived from data, computed as the mean-squared error *e*_*i*_(*t*) along all *i=1*, 2,…, *N* target neurons (e.g., Figures x,y,z); and 2) the proportion of variance explained (pVar) as one minus the ratio of the Frobenius norm of the difference between the neural data and outputs of the network compared to the variance of the data (e.g., see figures x,y,z):

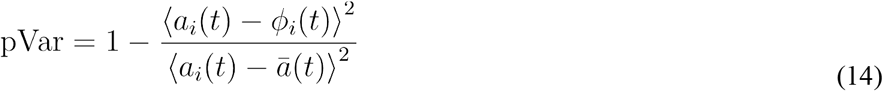

#### Analyzing the Directed Interaction matrix J after training

The directed interaction matrix inferred by the Model RNN quantifies the strength of interactions between the units in the network. These values can be either positive or negative, suggesting excitatory or inhibitory effects on the target neuron, respectively. Since the RNNs we build are extensively constrained by neural dynamics, we find that it is possible to consistently infer similarly distributed matrices, even after starting from different random initializations (e.g., **Figure S6c,h**). Thus the statistical properties of the interaction strengths we derive from data-constrained RNNs can be reliably compared across brain regions, as well as between RNNs trained to match a range of experimental datasets from different species. In this paper, we summarized the statistical properties of such model-derived interaction strengths by computing histograms using the total number of elements of either the full **J** matrix or specific submatrices containing the strength and type of interactions within and between individual brain regions. Notably, when analyzing these matrices, we scaled the distributions by the square root of the number of source units to account for differences in population sizes. We also normalized each histogram by the maximum value to facilitate comparison between matrices derived from RNNs of different sizes, and visualized the distributions using a logarithmic scale. These distributions could be further summarized and quantified by metrics such as the median, standard deviation, skewness, or kurtosis.

### Computing current sources to specific brain regions

The activity of each target unit in the Model RNN at each time step is computed as the product of the corresponding row of the directed interaction matrix and the activity of all source units at the previous time step. Thus, it follows that the current into the *i*th target unit, *I*_*i*_(*t*), can be estimated by multiplying the corresponding row of the directed interaction matrix with the activity of all of the source units.

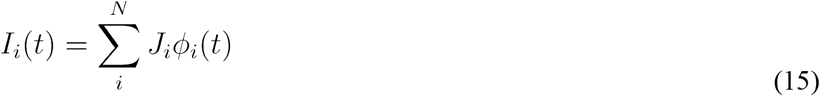

Since this is a linear operation, the above equation can be rewritten as a sum of separate contributions from each of source units:

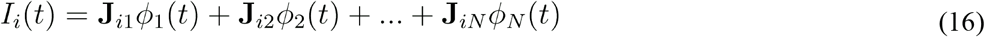

CURBD adopts this linear decomposition to study brain-wide currents between active neurons across multiple interacting brain regions. Based on Eq. 16, the total current input into a single target region from another source region can therefore be computed by grouping the currents from the source region weighted by the strength of the directed interactions between them. In this manuscript, we computed the currents in the target regions using the weights in different submatrices as described here. However, this method can be readily extended to separately infer excitatory (or inhibitory) currents by first setting all of the negative (or positive) values in the **J** matrix to be zero and then repeating the summation in Eq. 16.

Due to the large number of free parameters in the Model RNN, i.e., order *N*^*2*^ elements for RNNs with *N* units, the training algorithm does not necessarily infer the precise entries, element-by-element, in the directed interaction matrix, even when ground truth simulated data originated via low-rank or smoothed connectivity (for details, see Ref. ^17^). However, we find consistent and reliable estimates, i.e., recapitulating statistical properties of groups of weights in the directed interaction matrices. Furthermore, after training, the RNNs are able to produce highly consistent dynamics even when starting from different initial conditions. In practice, when taking the dot product of **J** and *ϕ*(*t*) to compute the currents for CURBD, random element-by-element fluctuations in the individual reconstructed weights between pairs of units are averaged out, but the overall population dynamics are preserved. For this reason, in its current state, CURBD is best applied to infer interactions between source and target brain regions with sufficient numbers of active neurons. Future extensions, e.g., those that incorporate known connectivity between regions^57,68^ or additional constraints from data, such as behavioral covariates, could provide reliable current estimates with finer granularity than at the level of individual regions and possibly across different behavioral states.

### Two-region model producing idealized, ground truth, simulated data to validate CURBD

#### Design of the generator model

We simulated a model that generated idealized ground truth data to test when CURBD approach would be the most effective at disentangling inter-region interactions, and to probe the conditions under which it would fail to perform optimally. We generated two 1000 unit RNNs, each with random connectivity weights drawn from a Gaussian distribution, as described in the initialization procedure for the Model RNN above. One RNN (corresponding to Region A) was driven by an external sinusoidal signal, *S*_*A*_(*t*), oscillating at *Π*/8 Hz, while the second (corresponding to Region B) was driven by another sinusoid, *S*_*B*_(*t*), oscillating at *Π*/3 Hz and phase shifted by *Π*/3. The two sinusoidal inputs began after two seconds of a simulated “resting state” during which the inputs to the RNNs were set to zero. These external inputs were connected to 33% of the units in their respective RNN with a fixed input weight, picked from a uniform distribution. The two RNNs were recurrently connected, with a varying percentage of neurons in each region (randomly selected) receiving inputs from the other region with a fixed weight of one. We computed the time-series activity of the *i*^th^ unit in the two RNNs, *r*_*A, i*_(*t*) *and r*_*B, i*_(*t*) for ten seconds of data using the following steps. We initialized the states of the two RNNs to random values between -1 and 1. For each subsequent time step, we computed the change in activity of each RNN unit *i* based on its inputs according to:

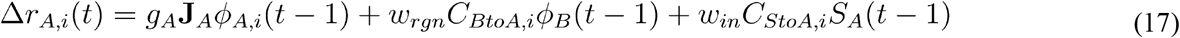

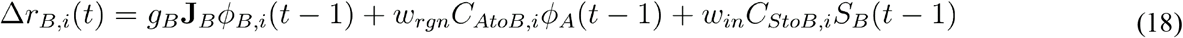

The scaling parameters *g*_*A*_ and *g*_*B*_ control how chaotic each RNN is, as described in the Model RNN training section above. *C*_*StoA*_ and *C*_*StoB*_ represent a binary connectivity vectors describing the connectivity of the external sinusoidal inputs to their respective regions. Similarly, *C*_*AtoB*_ and *C*_*BtoA*_ are binary vectors describing the connectivity between regions A and B. The fraction of entriess in the above inter-region connectivity vectors set to 1 is defined as *p*_*rgn*_. *w*_*rgn*_ and *w*_*in*_ are scalars that set the connection weights for the sinusoidal inputs and inter-region connections, respectively. The activity of each RNN unit *i* was then computed according to:

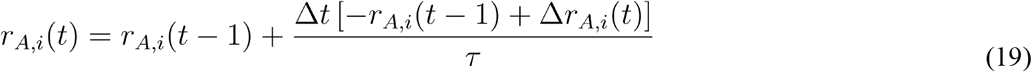

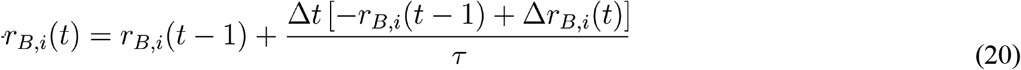

Lastly, the activity of each RNN unit *i* was transformed into a firing rate by passing through the nonlinearity, as described above:

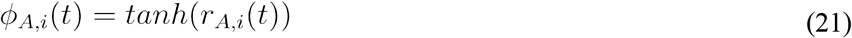

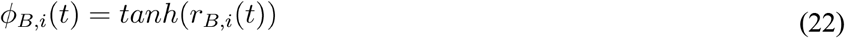

#### Checking robustness of CURBD over a range of simulation parameters for the Generator model

We repeated the ground truth simulations sweeping over a broad range of parameters applicable to the generator model (**Figure S3g**): 1) *g*_*A*_, the dynamical regime of Region A; 2) *w*_*rgn*_, the strength of the weights of the recurrent connections between the networks; and 3) *p*_*rgn*_, the proportion of neurons in Regions A and B receiving input from the other region. This parameter-sweeping process helped us explore how effectively CURBD operates to untangle currents resulting from the external sinusoidal inputs as the properties of the modeled networks change. The remaining parameters were fixed for all of these simulations. Values for all of the parameters we tested are provided in **Table 1**.

**Table 1.** Two-region generator model parameters.

| Parameter | Description | Value(s) | Parameter | Description | Value |
| --- | --- | --- | --- | --- | --- |
| $g_A$ | Region A chaos | [0.9, 1.0, 1.1, 1.3, 1.5, 1.8, 2.5] | $w_{rgn}$ | Inter-region weight | [0.001, 0.01, 0.05, 0.1, 0.25, 0.5, 1.0] |
| $g_B$ | Region B chaos | 1.5 | $p_{rgn}$ | Inter-region connection proportion | [0.01, 0.05, 0.1, 0.25, 0.5, 1.0] |
| $\tau$ | Time constant of model units | 0.1 | $w_{in}$ | External input weight | 1.0 |
| $T$ | Simulation time | 10 | $\Delta t$ | Simulation time step size | 0.01 |

For each combination of parameters in the generator model, we trained a 2000-unit Model RNN (**Figure S3b**) to reproduce the activity of the two Regions from the generator model using the algorithm described above. The parameters chosen for this “fit” Model RNN (**Table 2**) were held fixed to ensure we studied the effect *only* of the network properties as we swept parameters, not of variations in the Model RNN. For each of these Model RNNs, we applied CURBD to Region A to assess how effectively we could isolate the currents from the two external inputs (the sinusoidal input driving Region A and the sinusoidal input driving Region B) from the population. Since each external input was effectively one-dimensional, we first reduced each estimated population-wide source current to a single component using PCA. We then computed how accurately we could infer the external inputs by directly comparing the correlation coefficient (denoted by R^2^ in **Figure S3f-g**) between the leading PC of each source current and each of the two sinusoidal external inputs. We repeated this analysis for different combinations of parameters to assess which parameter regimes consistently gave the highest R^2^ values.

**Table 2.** Model RNN training parameters for all datasets.

| Parameter | Description | 2-region simulation | 3-region simulation | Mouse dataset | Monkey dataset | Human dataset |
| --- | --- | --- | --- | --- | --- | --- |
| $g$ | Chaos | 1.5 | 1.5 | 1.5 | 1.5 | 1.5 |
| $\tau$ | Time constant for model units | 0.1 | 0.1 | 0.3 | 0.01 | 0.0075 |
| $P_0$ | Learning rate | 1.0 | 1.0 | 1.0 | 1.0 | 1.0 |
| $\tau_{WN}$ | Time constant of filtered white noise inputs | 0.1 | 0.1 | 0.1 | 0.1 | 0.1 |
| $w_{WN}$ | White noise input weight | 0.01 | 0.01 | 0.001 | 0.0001 | 0.001 |
| $n_{iterations}$ | Number of training iterations | 100 | 500 | 2500 | 1500 | 2500 |
| $\Delta t$ | Data time step size (s) | 0.01 | 0.01 | 0.1866 | 0.01 | 0.01 |
| $\Delta t_{RNN}$ | Model RNN integration step size (s) | 0.001 | 0.001 | 0.0467 | 0.0005 | 0.001 |

### Three-region ground truth simulation to validate CURBD

#### Design of the generator model

We designed a second idealized ground truth simulation to validate whether CURBD could effectively infer source currents between different interacting regions even when there are no external inputs driving a particular neural population. We simulated three 1000-unit RNNs using a generator model similar to the two-region model described above (**Figure 3a**). However, rather than sinusoidal inputs as in the two-region ground truth model, two of the three interconnected RNNs received time-varying patterns of inputs from other model networks. The external inputs driving Region B were provided by a network generating a Gaussian “bump” propagating across the network sequentially, *SQ*(*t*). The sequence began 2 seconds after the start of the simulation and ended 4 seconds later, with each sequentially activating unit *i=1*, 2,…, *N* behaving according to:

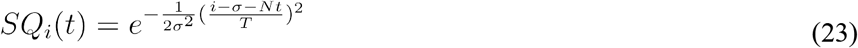

where *σ* denotes the width of the bump across the population (here, 20% of the units), *N* represents the population size (here, 1000 units), and *T* represents the total simulation time of twelve seconds.

The external input to Region C was provided by another 1000-unit network generating a fixed point, *FP*(*t*), for 8 seconds that instantaneously shifted to a new fixed point for an additional 4 seconds. The fixed points were generated by sampling *SQ*(*t*) at two different time points (*t=*2s and *t=*5s) and holding them at the sampled value of firing rate for the duration of the fixed point. The external inputs were connected to 50% of the units in their respective regions (randomly selected) with a fixed negative weight (inhibitory) for Region B and positive weight (excitatory) for Region C. The third RNN (Region A) received only the recurrent inputs from the other two RNNs, no external drive. The Region A RNN was modeled at a different value of *g* from the other two networks, yielding distinct dynamics (**Table 3**). The following update equations governed the interactions of the regions at each time step (with a resolution of *Δt=0*.*01*), with the subsequent activity evolving similarly to the two-region simulation described Eqs. 19-22:

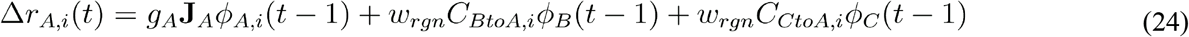

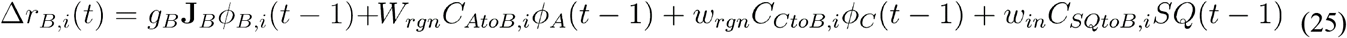

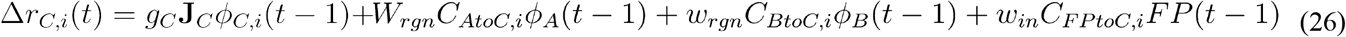

**Table 3.** Three-region generator model parameters.

| Parameter | Description | Value | Parameter | Description | Value |
| --- | --- | --- | --- | --- | --- |
| $g_A$ | Region A chaos | 1.8 | $w_{rgn}$ | Inter-region connection weight | 0.01 |
| $g_B$ | Region B chaos | 1.5 | $p_{rgn}$ | Fraction of inter-region connections | 0.01 |
| $g_C$ | Region C chaos | 1.5 | $w_{in}$ | External input weight | 1.0 |
| $\tau_{true}$ | True decay constant | 0.1 | $\sigma$ | Width of sequential and fixed point-bumps (number of RNN units) | 200 |
| $T$ | Simulation time | 12 | $\Delta t$ | Simulation time step | 0.01 |

#### Description of the Model RNN and CURBD analysis

We trained a 3000-unit Model RNN to match the activity of the three-region generator model using the procedure described above. We found that the Model RNN reproduced the simulated data accurately over a wide range of parameters; for the simulations reported in this paper (**Figures 3, S1, S2**), we used the values reported in **Table 2**. We performed CURBD to infer the nine source currents governing interactions between the three regions. We reduced the dimensionality of the full 1000-unit population of each of the three regions and the source currents using PCA. We chose the leading five dimensions for the following analyses, which sufficed to capture more than 95% of the total variance in each source current, though we observed similar results with other assumed dimensionalities (data not shown).

Since the true connectivity of the network was defined in the simulated dataset, we computed the ground truth currents between each region to isolate the effect of isolated inputs from the source regions on the target region, including how the input activity would propagate through the recurrent connections of the target region. We adapted the update equations defined above to compute the three current sources into one region at each time step, here using Region A as an example:

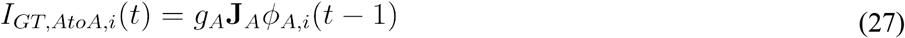

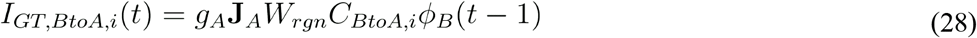

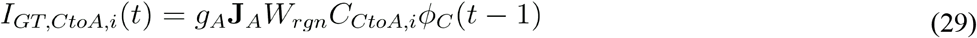

The same process was performed for Regions B and C using similar equations. We performed the same dimensionality reduction analysis on the ground truth currents as on the inferred source currents from CURBD. Since the Model RNN was trained to reproduce time-varying activity from all the units in the multi-region generator model, each inferred source current has the same dimensionality as the ground truth current, and is embedded within the same high-dimensional space of the population activity of the respective simulated region. We could thus directly compare each leading PC using VAF (**Figures 3g, S1**).

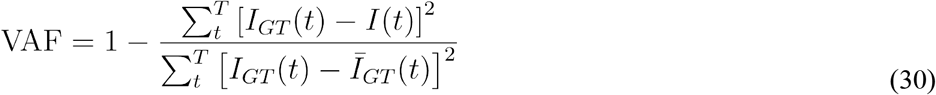

#### Comparison of inferred inter-region currents to shared dynamics identified by Canonical Correlation Analysis

We compared the performance of CURBD to an analogous decomposition obtained by canonical correlation analysis (CCA)^28^. CCA obtains an optimal linear transformation relating the dimensionality-reduced population activity of the source and target regions to identify shared dynamics. In brief, we first took the low-dimensional trajectories of each region and performed a QR decomposition to identify for each region of the resulting **Q**, which provides an orthonormal basis for the column space of the low dimensional trajectories. For any pair of regions, for example Region A and Region B, we performed a singular value decomposition of the inner product of the corresponding **Q** matrices:

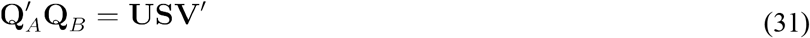

This process effectively finds new dimensions within the manifold of Region A (denoted by **U**) and Region B (denoted by **V**) that maximize the correlation between the two trajectories. To analyze the shared dynamics between the two regions, we projected the activity of either region onto the corresponding axes. Unlike CURBD, the mapping obtained from CCA (and similar methods of inferring functional connectivity only from the covariance matrix of recorded neural activity) is not directional and is purely correlational. Thus, only one “current” can be obtained for each pair of regions. We compared the VAF by the first component identified by CCA to the first PC of the ground truth currents to assess the effectiveness of this approach (**Figure S2**).

#### Addressing partial sampling issues present in experimental data

We repeated the above simulation to determine whether or not CURBD is effective when only a fraction of the total multi-regional activity is ‘observed’ by the Model RNN. This control analysis addresses partial sampling issues present in real data when activity can be experimentally measured from only a relatively small fraction of the total number of neurons in a region. To simulate this scenario and test the efficacy of CURBD in the face of partial sampling issues, we trained Model RNNs to match activity from 5%, 20%, 50%, and 100% of the available neurons in each region (randomly selected) of the ground truth multi-region generator model. We repeated the simulation ten times at each subsampling level to help account for variability in the random sampling of neurons, as well as variability from different random initializations of the **J** matrix. Such variability scales inversely with network size for Gaussian weights; the ten repetitions at 100% sampling thus provide a lower-bound on the variability that would be expected within this model. Unlike the initial Model RNN analysis where every neuron was sampled, in the subsampling case, we can no longer guarantee that the axes should be oriented similarly in PC space and VAF is not a reliable measure of how well the method performed. Thus, here we again employed CCA not to identify shared dynamics between regions, but to compensate for differences in the number of sampled neurons generating the dynamics of a single region^28^, In this application, CCA provides a quantitative “similarity index”–quantified by the canonical correlation of the leading aligned dimension–of the population dynamics between the currents identified by CURBD and the ground truth currents (**Figure 3H**).

### Multi-region calcium fluorescence recordings in mice

#### Surgery

All experimental procedures were approved by the Harvard Medical School Institutional Animal Care and Use Committee and were performed in compliance with the Guide for the Care and Use of Laboratory Animals. Two female mice expressing GCaMP6s (C57BL/6J-Tg(Thy1-GCaMP6s)GP4.3, The Jackson Laboratory, stock 024275) were implanted with cranial windows over the cortical surface. Mice were 3-5 months old at the time of surgery, and given an injection of dexamethasone (3 µg per g body weight) 4-8 h before the surgery. Mice were anesthetized with isoflurane (1-2% in air). A cranial window surgery was performed to either fit a ‘crystal skull’ curved window (LabMaker UG) exposing the dorsal surface of both cortical hemispheres^80^, or to fit a stack of custom laser-cut quartz glass coverslips (three coverslips with #1 thickness each (Electron Microscopy Sciences), cut to a ‘D’-shape with maximum dimensions of 5.5 mm medial-lateral and 7.7 mm anterior-posterior, and glued together with UV-curable optical adhesive (Norland Optics NOA 65), exposing the left cortical hemisphere. The dura was removed before sealing the window using dental cement (Parkell). A custom titanium headplate was affixed to the skull using dental cement mixed with carbon powder (Sigma-Aldrich) to prevent light contamination. A custom aluminum ring was affixed on top of the headplate using dental cement. During imaging, this ring interfaced with a black rubber balloon enclosing the microscope objective for light-shielding.

#### Imaging and behavior setup

Data were collected using a large field of view two-photon microscope assembled as described in Ref. ^31^. In brief, the system contained a combination of a fast resonant scan mirror and several large galvanometric scan mirrors allowing for especially large scan angles. Paired with a remote focusing unit to rapidly move the focus depth, this setup enabled random access imaging in a field of view of 5-mm diameter with 1 mm depth. The setup was assembled on a vertically mounted breadboard whose XYZ positions and rotation were controlled electronically via a gantry system (Thorlabs). Thus, to position the imaging objective with regards to the mouse, the position and rotation of the entire microscope were adjusted while the position of the mouse remained fixed. Mice were head-fixed and placed on an air-suspended 8-inch diameter Styrofoam spherical treadmill that enabled spontaneous running. Using two optical sensors (ADNS-9800, Avago Technologies), we tracked the treadmill velocity, which was translated into pitch, roll, and yaw velocity using custom code on a Teensy microcontroller (PJCR) as a readout of the mouse’s running speed and direction. Individual recording sessions lasted from 45–60 minutes. Mice were extensively acclimated to head-fixation and running on the treadmill before data collection. We recorded behavioral and neural activity while mice spontaneously ran on the ball. The room was kept in complete darkness throughout the experiment. We defined running bouts as periods when the ball movement speed crossed a fixed threshold set to be the 90th percentile of the running speeds throughout the session.

#### Image acquisition

The excitation wavelength was 920 nm, and the average power at the sample was 60-70 mW. The microscope was controlled by ScanImage 2016 (Vidrio Technologies). We targeted four distinct regions in the left cortical hemisphere: primary visual cortex (V1), secondary motor cortex (M2), posterior parietal cortex (PPC), and retrosplenial cortex (RSC). These regions were targeted based on retinotopic mapping (see below). In each region, we acquired images in layer 2/3 from two planes spaced 50 µm in depth, at 5.36 Hz per plane at a resolution of 512 x 512 pixels (600 µm x 600 µm).

#### Retinotopic mapping for selecting Ca2+ imaging locations

We performed retinotopic mapping in the mice used for calcium imaging experiments as previously described in Ref. ^63^. Mice were lightly anesthetized with isoflurane (0.7–1.2% in air). GCaMP fluorescence was imaged using a tandem-lens macroscope where excitation light (455 nm LED, Thorlabs) was filtered (469 nm with 35 nm bandwidth, Thorlabs) and reflected onto the brain through a camera lens (NIKKOR AI-S FX 50 mm f/1.2, Nikon) focused 400 μm below the brain surface. GCaMP emission light was collected using the same lens, filtered (525 nm with 39 nm bandwidth, Thorlabs), and imaged with another camera lens (SY85MAE-N 85 mm F1.4, Samyang) and a CMOS camera at 60 Hz (ace acA1920-155um, Basler). These images were synchronized to visual stimuli presented on a gamma-corrected 27 inch IPS LCD monitor (MG279Q, Asus). The monitor was centered in front of the mouse’s right eye at an angle of 30 degrees from the mouse’s midline. The visual stimulus, a spherically corrected black and white checkered moving bar^81^ (12.5 degree width, 10 deg/s speed), was presented in 7 blocks, each consisting of 10 repeats of 4 movement directions (up, down, forward, backward). To produce retinotopic maps, we calculated the temporal Fourier transform at each pixel of the imaging data and extracted the phase at the stimulus frequency^82^. These phase images were averaged across repetitions for a given movement direction and smoothed with a Gaussian filter (25 μm s.d.). Lastly, we calculated field sign maps by computing the sine of the angle between the gradients of the average horizontal and vertical retinotopic maps.

For each retinotopic mapping session, we also acquired an image of the superficial brain vasculature pattern under the same field of view. We then acquired a similar brain vasculature image under the large field of view two-photon microscope. These two reference images were manually aligned and used to directly locate V1 and PPC locations for two-photon imaging. The location for RSC imaging was positioned adjacent to the midline and about 300 μm anterior of the PPC location. The location for M2 imaging was positioned one millimeter anterior of the RSC location.

#### Pre-processing of imaging data

We used custom code to correct for motion artifacts, as described in Ref. ^83^. In brief, motion correction was implemented as a sum of shifts on three distinct temporal scales: sub-frame, full-frame, and minutes- to hour-long warping. After motion correction, ROIs were extracted using Suite2P^84^. Afterwards, somatic sources were identified with a custom two-layer convolutional network in MATLAB trained on manually annotated labels to classify ROIs as neural somata, processes, or other^83^. Only somatic sources were used. This yielded large populations from neurons from each of the four targeted regions in Mouse A and Mouse B (**Table 4**).

**Table 4.** Simultaneous neuron yield for the mouse dataset.

| Brain Region | Mouse A | Mouse B |  |
| --- | --- | --- | --- |
|  |  | Session 1 | Session 2 |
| V1 | 385 | 114 | 176 |
| M2 | 208 | 57 | 118 |
| PPC | 1124 | 247 | 462 |
| RSC | 1070 | 581 | 688 |
| *Total* | 2787 | 999 | 1444 |

After identifying individual neurons, we computed average fluorescence in each ROI and converted this value into a normalized change in fluorescence (*ΔF/F*). We corrected the numerator of the *ΔF/F* calculation for neuropil by subtracting a scaled version of the neuropil signal estimated per neuron during source extraction:

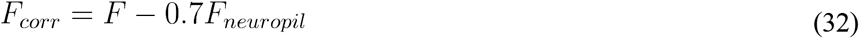

We estimated the baseline fluorescence (*F*_*base*_) of this trace as the 8th percentile of fluorescence within a 60-s window and subtracted this baseline to get the numerator:

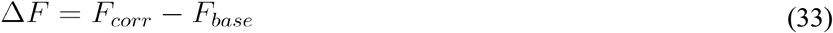

We divided this by the baseline (again 8th percentile of 60s window) of the raw fluorescence signal to get *ΔF/F*. We deconvolved the *ΔF/F* trace per neuron using the constrained AR-1 OASIS method44. We initialized the decay constants at two seconds and then optimized separately for each neuron. To fit the Model RNN, we temporally smoothed the sparse deconvolved spike estimates using a Gaussian kernel with four times the width of the sampling rate. We applied the same filter to the behavioral signals (pitch, roll, and yaw of the ball) to preserve the temporal relationship with the neural activity. For visualization of the neural population activity in the heatmaps of **Figures 4 and S4**, we scaled each neuron by the mean of the total activity.

#### CURBD analysis to infer source currents from mouse data

For each mouse, we trained Model RNNs (Mouse A: 2787 units; Mouse B, Session 1: 999 units; Mouse B, Session 2: 1444 units) to match the time-series Ca^2+^ data from the four regions. We used identical parameters for each Model RNN (**Table 2**).

We applied CURBD to infer the sixteen source currents comprising the multi-region population activity. We first assessed how much unique explanatory power each source current had in the total V1 population. We developed a partial coefficient of determination analysis to quantify this as follows. We subtracted each source current one-by-one from each V1 neuron and computed the sum-squared error of this difference and the recorded neural data. We then computed the sum-squared error of the full Model RNN fit compared to the recorded neural data. We defined the unique variance explained by the source current according to:

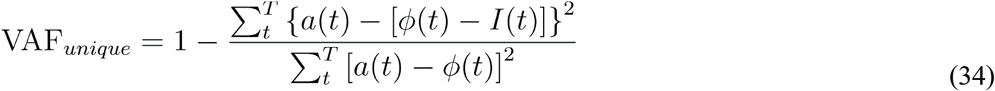

where *I(t)* denotes the source current that is being evaluated. Effectively, this computes the variance that cannot be explained by any of the three remaining source currents. Importantly, this metric can be computed at individual time points. We normalized each calculation by the sum of the four unique variances at each time point to give a proportion of unique variance explained by each source current. For cleaner presentation, we smoothed these normalized traces with a Gaussian kernel of width 500 ms (**Figure 4h**). We then reduced the dimensionality of all sixteen source currents using PCA, selecting a 5-dimensional manifold which sufficed to explain more than 80% of the total variance in all source currents. We trained Wiener cascade filters, a type of linear-nonlinear decoder^85^, to predict the running speed using the five-dimensional activity of each source current at each time step as well as the most recent 5 time steps of history. To perform cross validation, we randomly withheld 20% of time steps (the test set) and trained the decoders using the remaining 80% of the data. We quantified the performance of each decoder output on the left-out test set of time steps using VAF, as described above. We repeated this process for 100 iterations, randomly leaving out 20% of time steps for the test set on each iteration, and averaged across all iterations for the final decoder performance (**Figure 4k-l**).

### Multi-region electrophysiology recordings in monkeys

#### Behavioral task

All procedures were reviewed and approved by the Icahn School of Medicine Animal Care and Use Committee. For detailed descriptions of the experimental setup and protocol, see Ref. ^38^, where these data were previously reported. In brief, two rhesus macaque monkeys (*Macaca mulatta*; Monkey D: female, 5.6 kg; Monkey H: Male, 11.0 kg) were trained to sit in a custom primate chair with their head restrained and fixate on a computer monitor for four seconds, before performing a Pavlovian conditioning task for liquid rewards. They fixated on a neutral gray square for 800-1000ms. They were then presented with one of three visual conditioned stimuli for 500-600 ms on each trial corresponding to three different reward outcomes: no reward (CS-), water (0.5 mL), and juice (0.5 mL). An additional trial type occurred with equal frequency in which no conditioned stimuli was presented, and the gray square persisted throughout the trial. On all trials, a small (0.1 mL) water reward was given two seconds after the stimulus onset. Conditioned stimuli varied between monkeys and consisted of gray shapes, covering 1.10° of visual angle for Monkey D and 2.45° for Monkey H. We trained the Model RNNs described below using all four trial types to utilize as much training data as possible, though in this paper we only analyzed the CURBD output for the three stimuli.

#### Surgical procedures and neural recordings

After training, each monkey was implanted with a titanium head restraint device followed by a plastic recording chamber over the exposed cranium of the left frontal lobe. During the behavioral experiments, tungsten microelectrodes (FHC, Inc. or Alpha Omega, 0.5-1.5 M at 1 KHz) or 16-channel multi-contact linear arrays (Neuronexus Vector array) advanced by an 8-channel micromanipulator (NAN instruments, Nazareth, Israel) were attached to the recording chamber and inserted into the brain. The targeted brain regions were located using stereotaxic coordinates and verified by T1-weighted MRI imaging with the electrodes implanted. Recordings from subcallosal ACC were made on the medial surface of the brain ventral corpus callosum. Amygdala recordings were made between 22 and 18.5 mm anterior to the interaural plane. Rostromedial striatum recordings were made in the anterior medial segment corresponding to the zone where subcallosal and basal amygdala projections overlap. Spikes from putative single neurons were captured online using a Plexon Multichannel Acquisition Processor and later isolated with Plexon Offline Sorter. The small number of neurons recorded in each experimental session were then pooled into a pseudopopulation. First, the spike trains for each neuron on each trial were converted to an estimated firing rate. The firing rates were aligned on the stimulus presentation for each trial, then averaged across all trials of each stimulus type for that session in each monkey to give substantially large pseudopopulations (**Table 5**). As with the mouse dataset, neural activity was visualized with a heatmap after scaling each neuron’s firing rate by its mean activity (**Figures 5b, S6**).

**Table 5.** Pseudopopulation sizes for the monkey dataset.

| Brain Region | Monkey D | Monkey H |
| --- | --- | --- |
| Amygdala | 128 | 66 |
| Subcallosal ACC | 112 | 50 |
| Striatum | 103 | 83 |
| *Total* | 343 | 199 |

#### CURBD analysis of monkey dataset

For each monkey, we trained Model RNNs (Monkey D: 343 units; Monkey H: 199 units) to match the pseudopopulation data for all four conditions. We used identical parameters for the Model RNNs for both monkeys (**Table 2**). In the previous simulations and the mouse dataset, the Model RNN learned a single dynamical system that reproduces the neural data based on one initial condition. However, here we have four different initial conditions corresponding to the four trial types, and we seek to learn a single dynamical system that reproduces all of them. To achieve this, we concatenated the time-series data from the four conditions and reset the state of the Model RNN to match the real neural data at the first time point of each new condition. We repeated each Model RNN fit an additional four times, yielding five runs in total, each starting from a different randomly initialized matrix ***J***_***0***_ each time. We performed CURBD using each Model RNN to infer the nine source currents comprising the full multi-region activity. We then quantified the magnitude of current arriving to each source region from each target region by summing the absolute value of the source currents at each time step. We averaged this across the five runs in each monkey to assess the consistency of our solutions (**Figure 5f-g**).

We performed systematic subsampling analyses to assess whether applying CURBD to pseudopopulation data would be reliable even if different numbers and types of neurons were recorded experimentally. We randomly subsampled between 50% and 90% of the available neurons to create new, smaller pseudopopulations for each monkey. We used CCA (similar to the description in X above) to compute a similarity metric between the currents inferred by CURBD from each subsampled population and the currents originally inferred by CURBD using all of the available neurons for each monkey (**Figure S7**).

### Multi-region electrophysiology recordings in humans

#### Behavioral task

The institutional review boards of Cedars-Sinai Medical Center and the California Institute of Technology approved all protocols. Detailed descriptions of the experimental procedures are described in ^40^, where these data were previously reported. In brief, we recorded from 13 adult participants being evaluated for surgical treatment of drug-resistant epilepsy that provided informed consent and volunteered for this study. Of these thirteen participants, eight did not have a sufficiently large number of neurons to create a population for CURBD and were thus excluded. Our final analyses focused on five participants (P44, P51, P56, P57, and P58 from the original manuscript). The participants were seated in a chair facing a screen and reported decisions using either button presses or eye movements. They each performed eight forty-trial blocks that alternated between two tasks. In the categorization task, participants classified pseudorandomly presented images as belonging to one of four target categories (human faces, monkey faces, fruits, or cars) with a “yes” or “no” response. In the memory task, participants were shown an image and asked “Have you seen this image before, yes or no?” to which they responded “yes” or “no”. In the first block, all images were necessarily novel (40 unique images). In all subsequent blocks, the participants viewed 20 new images that were randomly intermixed with 20 familiar images. The 20 repeated images remained the same throughout the remainder of the session. We ignored trials where the participant provided an incorrect response (e.g. mistakenly identifying a novel image as familiar). This gave sixteen different conditions: four blocks of trials for each of two different tasks, each with correct “yes” and “no” responses. Participant task performance tended to improve throughout the session. Thus, we primarily focused our analysis on blocks 4, 6, and 8 (the final three memory blocks) when performance was highest.

#### Neural recordings

As participants performed this task, we recorded bilaterally from the amygdala, hippocampus, pre-supplementary motor area (preSMA), and dorsal anterior cingulate cortex (dACC) of each participant using microwires embedded in a hybrid electrode^41^. Electrode locations were confirmed using post-operative MRI or CT scans. We identified putative single neurons using a semi-automated spike sorting procedure. For the purposes of the CURBD analyses, we pooled recordings from each region from either hemisphere to ensure that we had sufficiently large populations for the dimensionality-reduction analysis. We estimated the instantaneous firing rate of each recorded neuron by convolving the spike train with a Gaussian kernel of width 150 ms. The relatively large width was necessary to accurately estimate firing rates for many low-firing neurons in the hippocampus and amygdala, and we opted to use a uniform width for all neurons. We created pseudopopulations by aligning each trial on the time of stimulus presentation and averaging across all trials for each task and correct response type (**Table 6**). In each trial, we kept two seconds before the stimulus presentation and three second after the stimulus presentation. As with the previous datasets, neural activity was visualized with a heatmap after scaling each neuron’s activity by its mean (**Figures 6f, S8**).

**Table 6.** Pseudopopulation sizes for the human dataset, reported as: Left Hemisphere / Right Hemisphere (Total).

| **Brain Region** | **P44**<br>(two sessions) | **P51**<br>(five sessions) | **P56**<br>(three sessions) | **P57**<br>(three sessions) | **P58**<br>(three sessions) |
| --- | --- | --- | --- | --- | --- |
| Hippocampus | 0 / 15 (15) | 25 / 38 (63) | 5 / 0 (5) | 17 / 0 (17) | 5 / 0 (5) |
| Amygdala | 3 / 1 (4) | 2 / 22 (24) | 33 / 43 (76) | 36 / 34 (70) | 54 / 61 (115) |
| preSMA | 62 / 0 (62) | 16 / 7 (23) | 8 / 14 (22) | 48 / 0 (48) | 65 / 72 (137) |
| dACC | 37 / 0 (37) | 72 / 17 (89) | 0 / 9 (9) | 43 / 0 (43) | 7 / 59 (66) |
| *Total* | 102 / 16 (118) | 115 / 84 (199) | 46 / 66 (112) | 144 / 34 (178) | 131 / 192 (323) |

#### CURBD analysis of human dataset

We trained Model RNNs based on the spiking pseudopopulation data from all sixteen conditions for each participant. As with the monkeys, we compensated for the discontinuities at trial boundaries by resetting the state of the Model RNN at the start of each condition. We used the same Model RNN parameters for all participants to ensure consistency (**Table 2**). We applied CURBD to infer the nine source currents comprising the multi-region interactions in the dataset. We reduced the dimensionality of each source current, as well as the full population activity of each region, using PCA. Since the stimulus was presented two seconds after the start of the trials, we defined the first two seconds to be the ‘resting state’ (**Figure 6g**). We then computed the Mahalanobis distance of each source current at each time step. This gave a time-varying estimate of how much the population activity or source currents responded to the stimulus. For each participant, we averaged across all five training runs to obtain the most reliable estimate of the current dynamics. We then averaged across all participants to show the group effect (**Figure 6h**).

## SUPPLEMENTAL FIGURES

**Figure S1.**
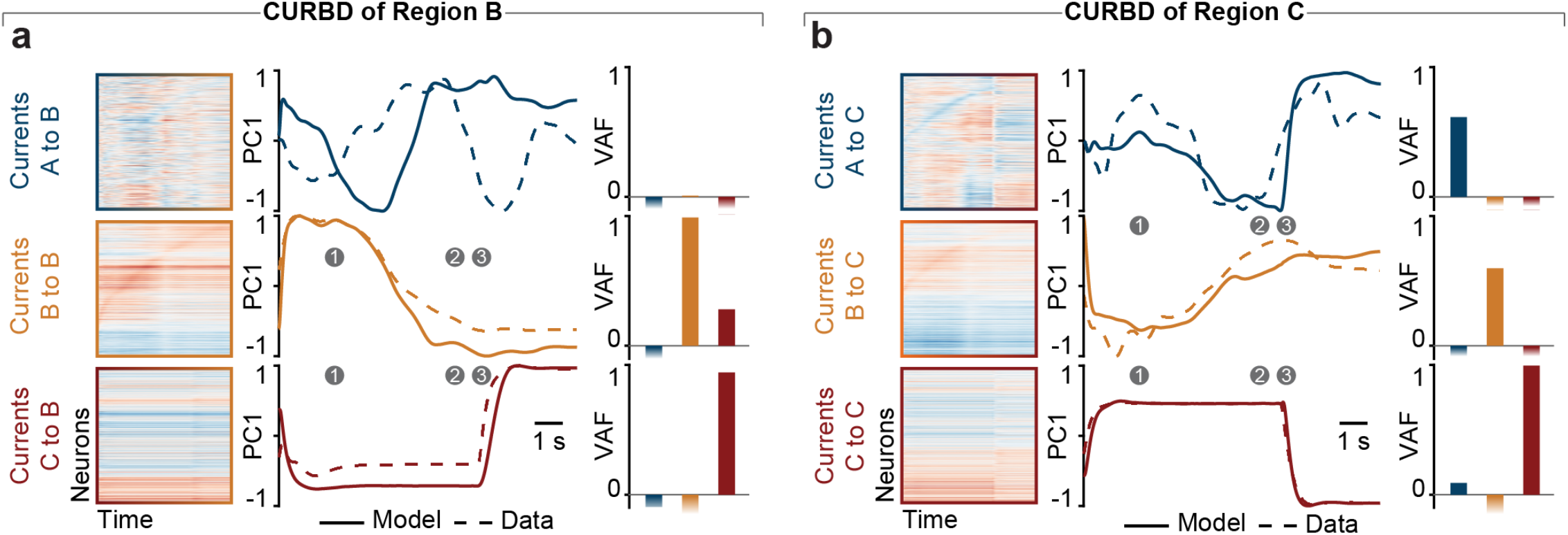
Supporting data for three-region ground truth simulation. **(a)** Analysis of source currents within Region B. Data presented as in Figure 3d. Currents from Region B and Region C are accurately reconstructed, though currents from Region A are missed, presumably due to the lack of strong external drive to Region A and the similar intrinsic dynamics between the three regions (VAF_AtoB_<0; VAF_BtoB_=0.98; VAF_CtoB_=0.94). **(b)** Analysis of source currents within Region C. Data presented as in Figure 3d. All three source currents are accurately reconstructed (VAF_AtoC_=0.61; VAF_BtoC_=0.60; VAF_CtoC_=0.99).

**Figure S2.**
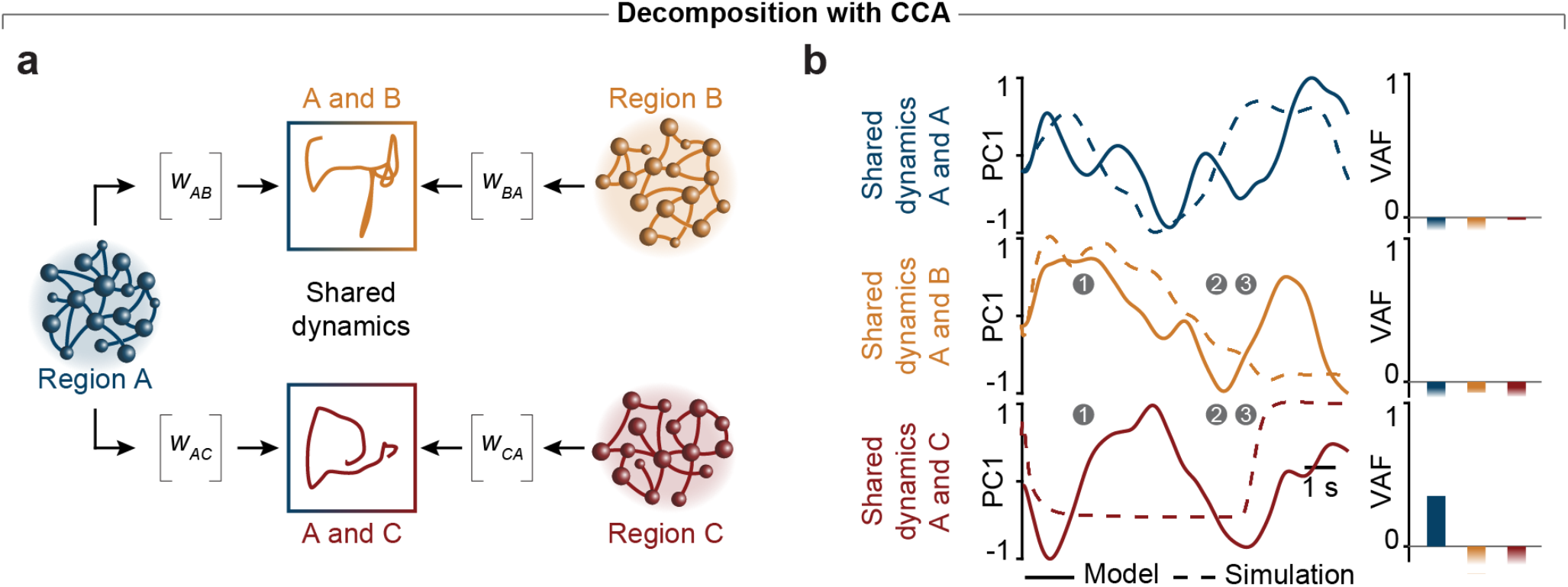
Decomposition of ground truth simulation using canonical correlation analysis. **(a)** CCA finds a single space capturing shared dynamics between each region, with a linear transformation (provided by the weight matrices *w*) relating each source and target region. However it does not provide a directional estimate of interactions. The shared dynamics plots show the aligned trajectories between pairs of regions projected onto the leading two aligned components. **(b)** Comparison of ground truth current inputs and shared dynamics identified by CCA. Unlike CURBD, the shared dynamics identified by CCA do not accurately match the ground truth current dynamics (VAF_AandA_<0; VAF_BandA_<0; VAF_CandA_<0).

**Figure S3.**
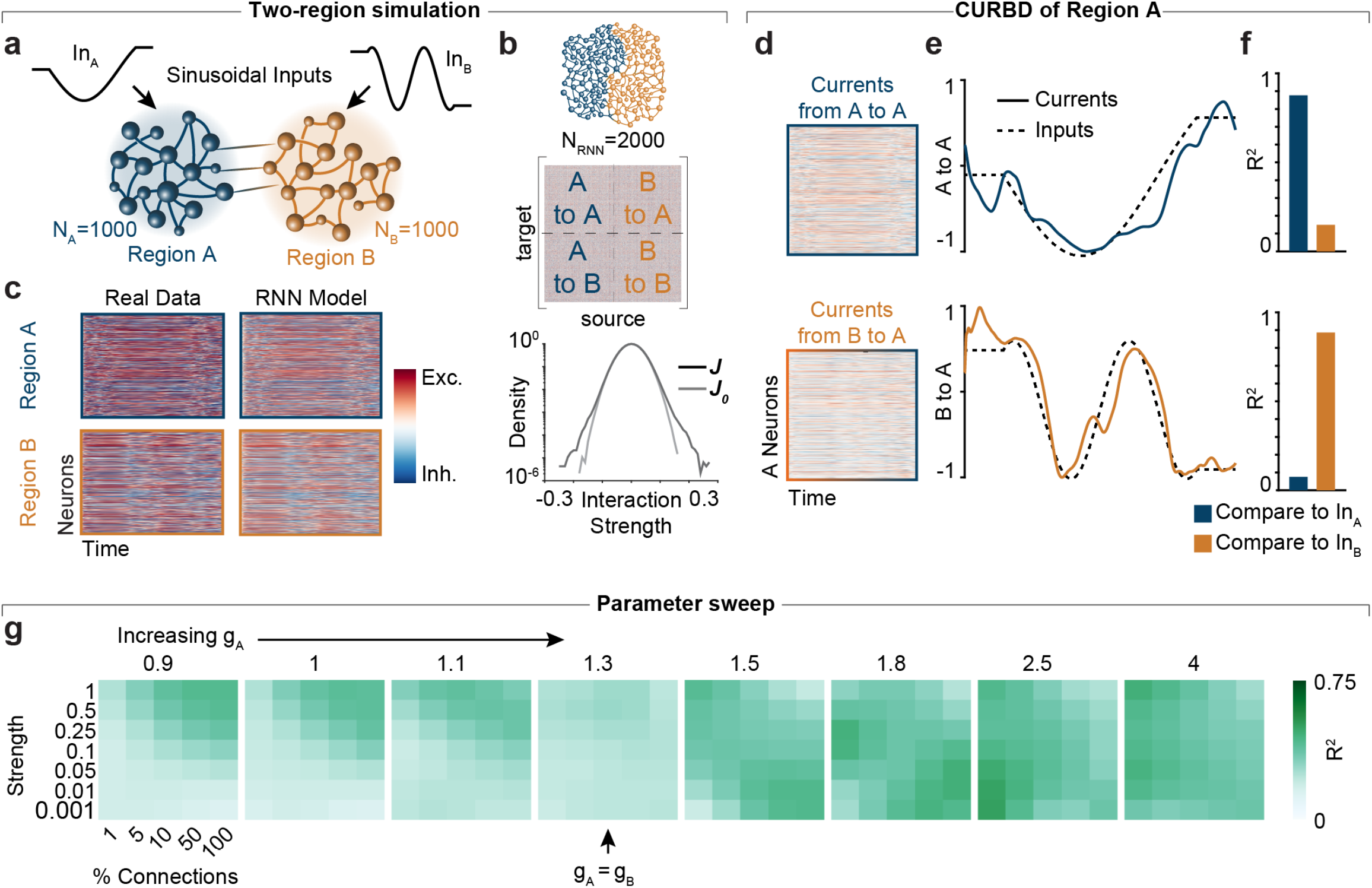
CURBD separates external inputs driving two interacting regions within specific dynamical regimes. **(a)** We simulated two interconnected RNNs representing distinct brain regions. Each was driven by a sinusoid of different frequencies. **(b-c)** We fit a Model RNN directly to the time-series data of the two regions to perform CURBD. From the Model RNN we obtained a matrix describing the directed interactions within and between each of the two regions. **(d)** We applied CURBD to obtain the currents driving each neuron in Region A from other Region A neurons (top) and from region B (bottom). **(e)** We performed PCA to identify the dominant component of each source current. The currents from Region A resembled the low-frequency sinusoid driving Region A, while the currents from Region B matched the higher-frequency sinusoid driving Region B. **(f)** We computed R^2^ values comparing the first PC of each source current to the two sinusoidal inputs. **(g)** Reconstruction accuracy of B to A currents for different simulation parameter values. We explored three key simulation parameters: i) the amount of chaos (g parameter; see Methods) from overdamped (g<1) to strongly chaotic (g>1.5); ii) the strength of the external inputs driving the system from very weak (0.001) to very strong (1); iii) the sparsity of inter-region connections from very sparse (1%) to full-rank (100%). Each heatmap shows the strength of inter-region connections against the percent of neurons receiving inter-region connections, and heatmaps going left to right show increasing g_A_. For low values of g_A_ corresponding to damped dynamics, the inputs can only be reconstructed with strong connectivity. When both regions have similar dynamics (g_A_ = 1.3) the currents cannot be accurately demixed. The optimal regime occurs when g_A_ and g_B_ have different dynamics, with a tradeoff between sparsity and strength of the inter-region connectivity.

**Figure S4.**
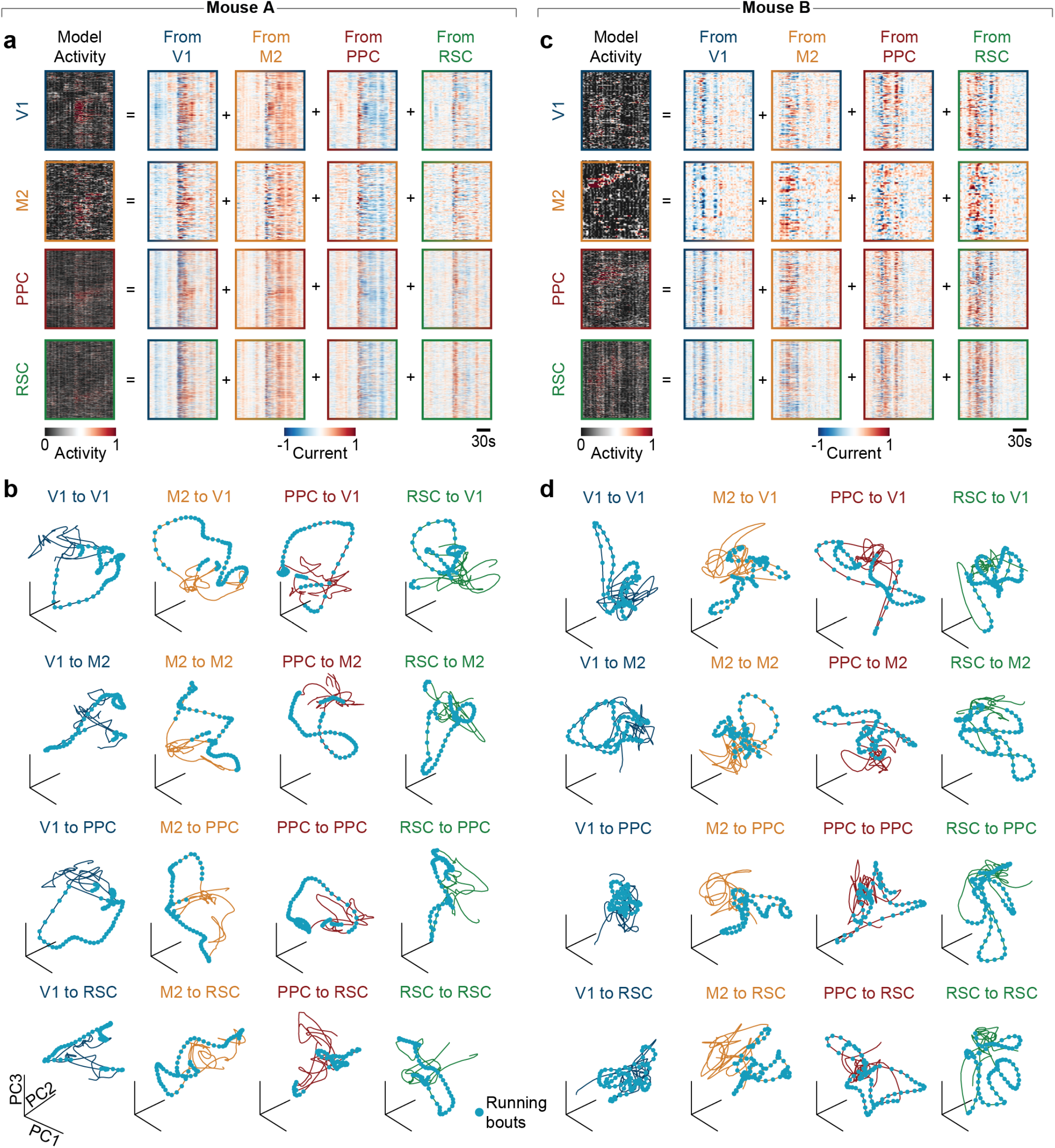
Supporting data for the multi-region mouse dataset. **(a)** Model RNN activity and source current activity for Mouse A. Figures reproduced from Figure 4g. **(b)** Current trajectories in the first three PCs for all sixteen source currents from Mouse A. The V1 source currents (top row) are reproduced from Figure 4h. **(c)** Data presented as in Panel a for Mouse B. **(d)** Data presented as in Panel b for Mouse B.

**Figure S5.**
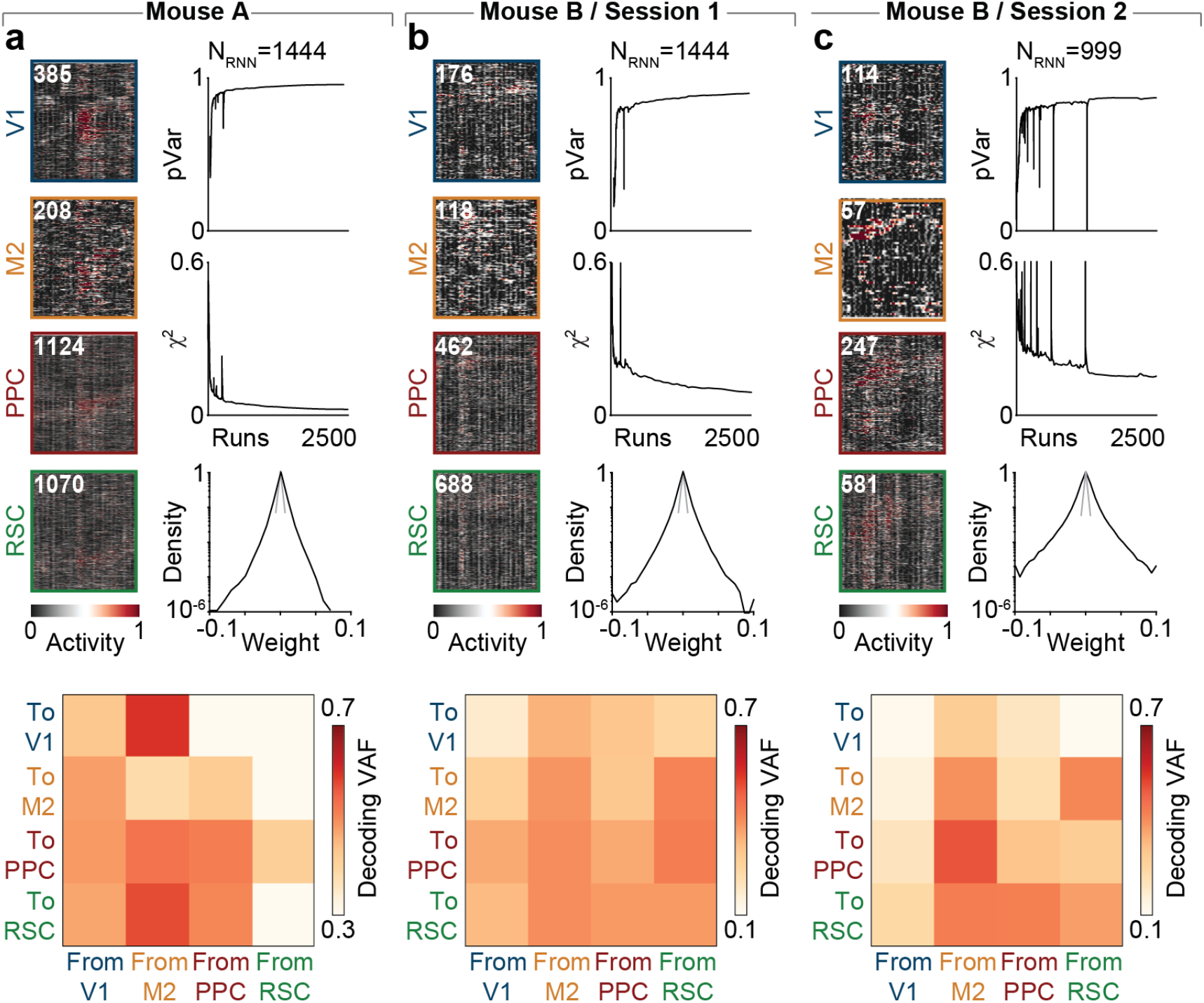
All recording sessions for the mouse dataset. **(a)** (Top) Model RNN output (left) and training performance (right) for the session from Mouse A. (Bottom) Decoding performance for the sixteen source currents. All data are reproduced from Figure 4. **(b)** (Top) Model RNN output (left) and training performance (right) for Session 1 from Mouse B. (Bottom) Decoding performance for the sixteen source currents. Portions are reproduced from Figure 4. **(c)** (Top) Model RNN output (left) and training performance (right) for Session 2 from Mouse B. (Bottom) Decoding performance for the sixteen source currents.

**Figure S6.**
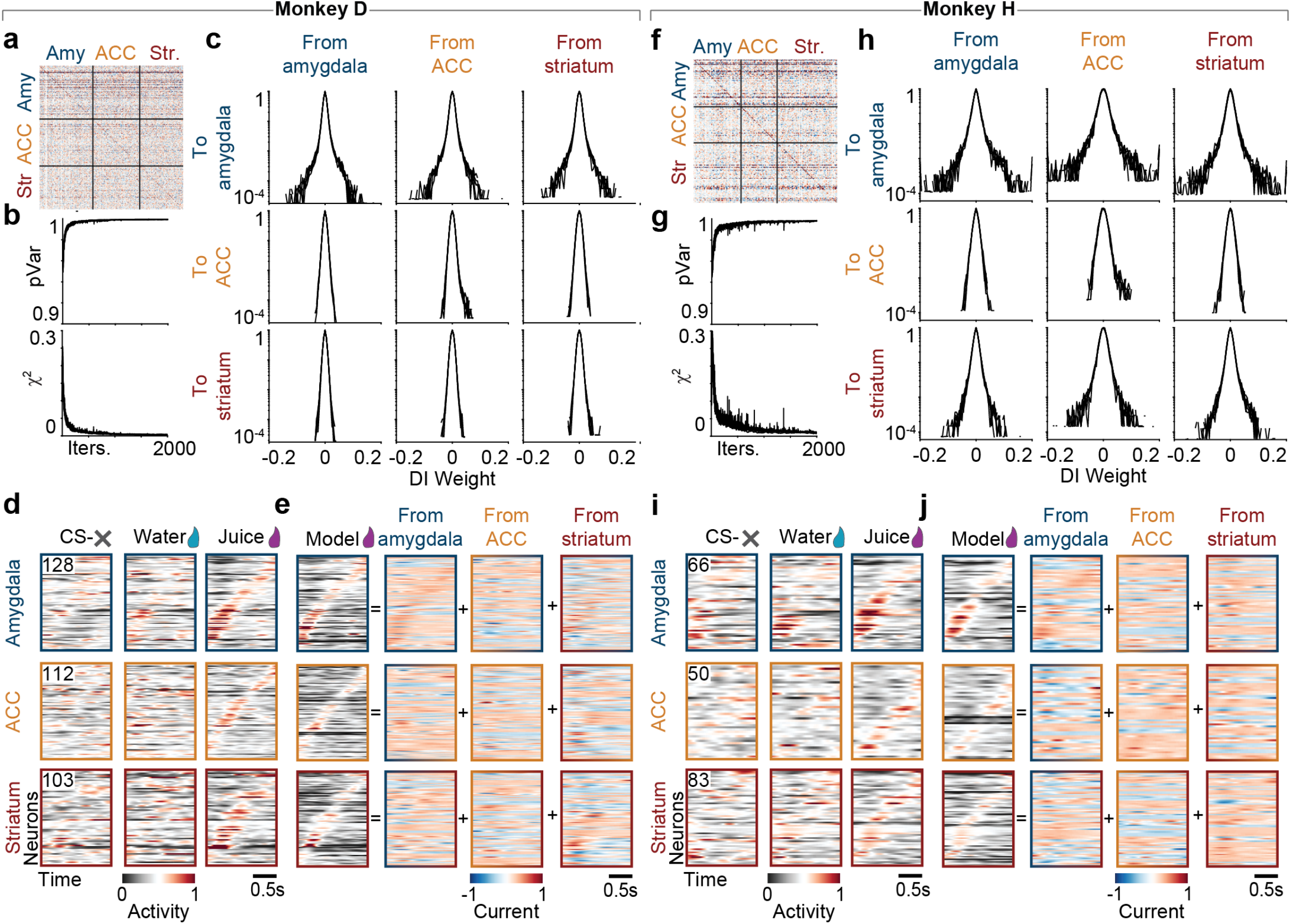
Supporting data for the multi-region macaque electrophysiology dataset. **(a)** Connectivity matrix for an example Model RNN fit to data from Monkey D. Neurons are ordered by region, starting with amygdala (Amy, blue), subcallosal anterior cingulate cortex (ACC, yellow), and rostromedial striatum (Str, red). **(b)** (Top) Proportion of variance explained (pVar) in the neural population as a function of training runs. Training results for five different random initializations are plotted to highlight consistency. (Bottom) Model error (*χ*^2^) for the five initializations shown above. **(c)** Distribution of weights in each submatrix used for CURBD. Each column corresponds to a source region, and each row to a target region. All five initializations are plotted to illustrate consistency. **(d)** Trial-averaged firing rates for the amygdala (top), subcallosal ACC (middle), and striatum (bottom) comprising the pseudopopulation dataset for Monkey D. Left plot shows data from the unconditioned stimulus (left), water stimulus (middle), and juice stimulus (right). All trials are aligned on presentation of the stimulus. Neurons in each region are sorted according to their time of peak activity in the Juice condition. **(e)** CURBD decomposition of activity in each region for the Juice trials. Left plots show the full Model RNN activity. The remaining plots show the inferred source currents to each target region (rows) from all source regions (columns). **(f-j)** Data for Monkey H presented as in Panels a-e.

**Figure S7.**
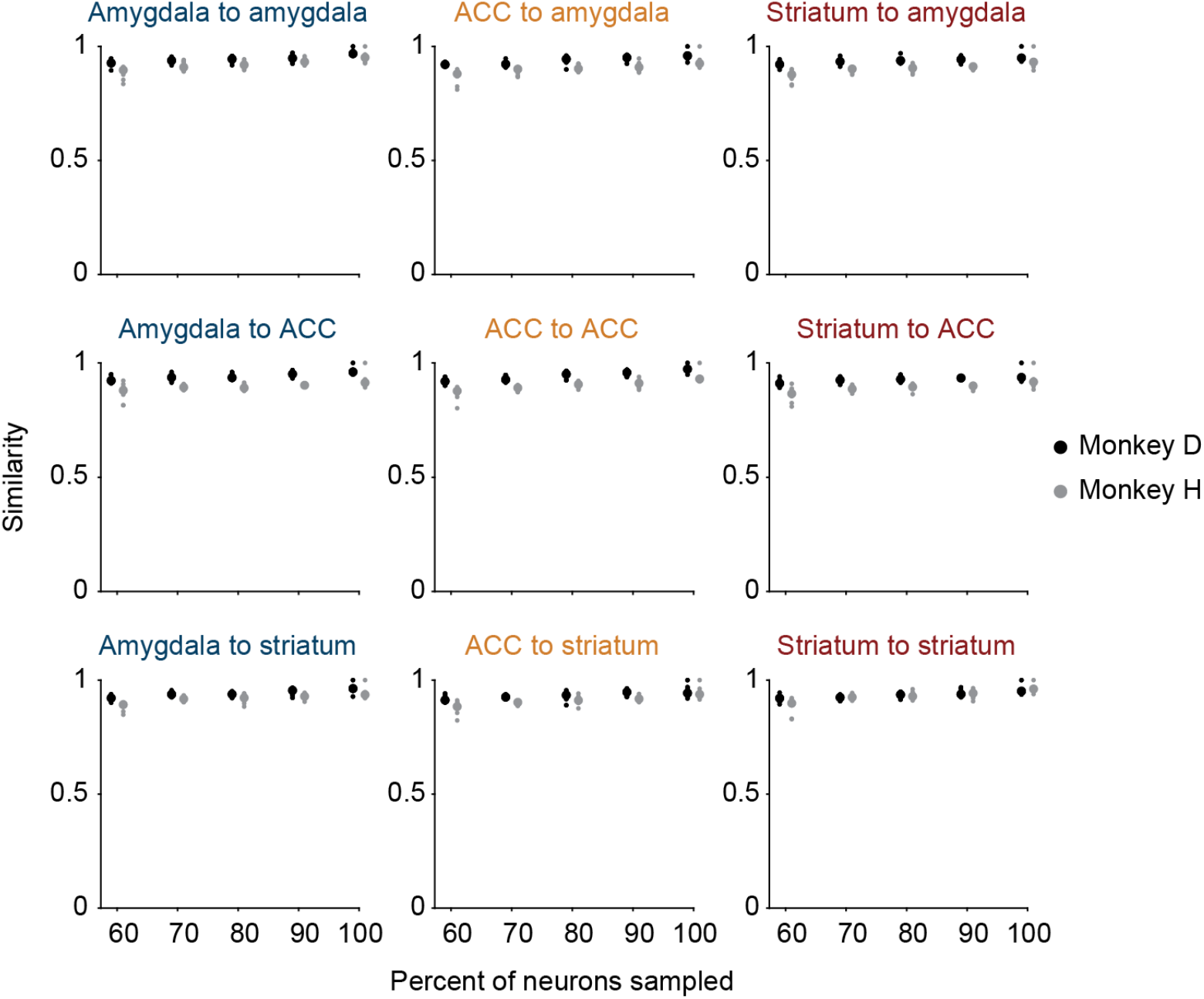
Consistent identification of current dynamics with random subsamples of recorded neurons. Mean canonical correlation in a twenty-dimensional space identified by PCA for each source current in Monkey D (black) and Monkey H (gray). Small dots indicate the results of ten random subsamples of the total neural population at each percentage level. Large circles indicate the median across iterations.

**Figure S8.**
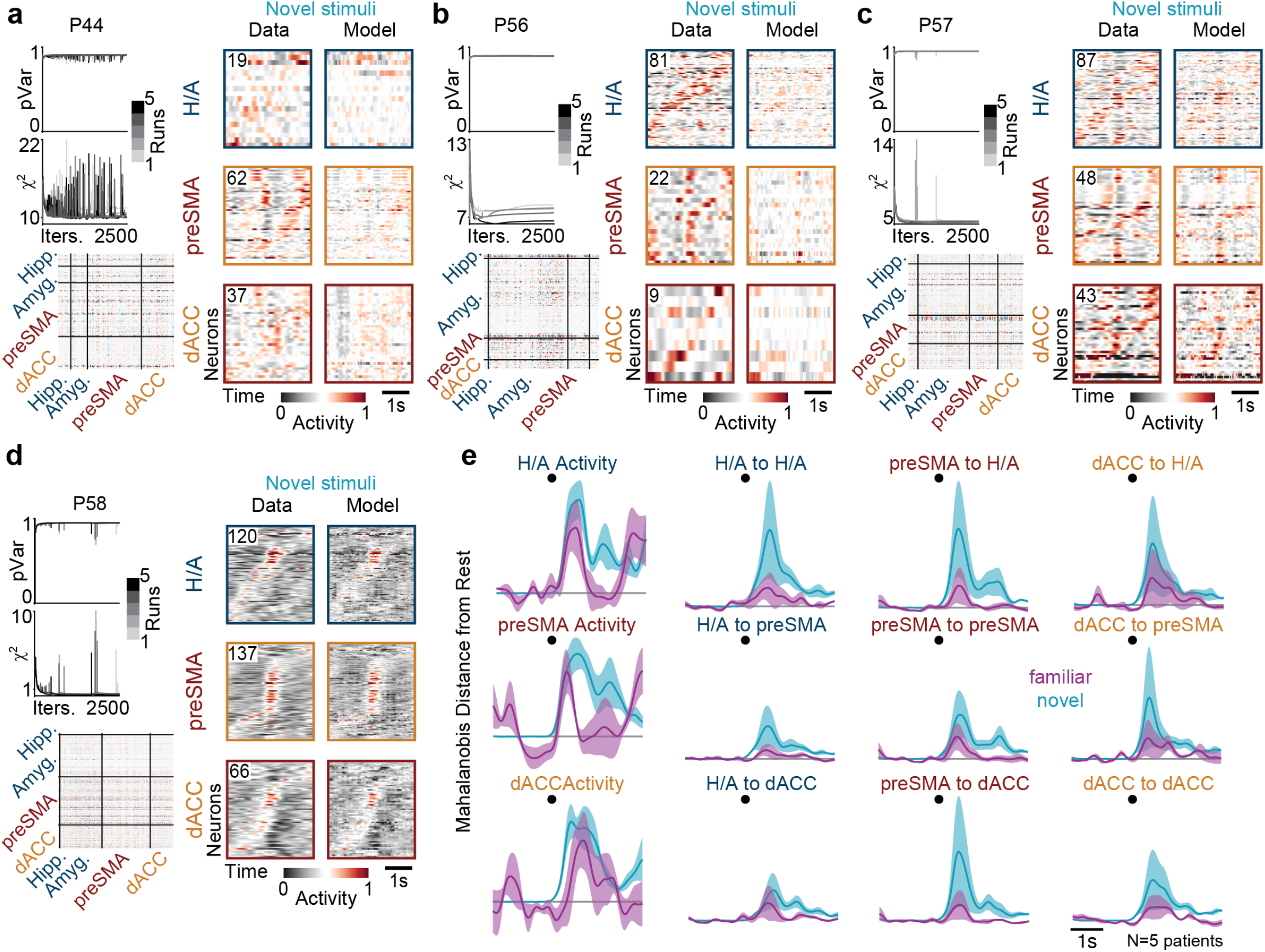
Supporting data for the multi-region human electrophysiology dataset. **(a)** Model RNN summary for P44. (Top left) Model RNN training performance (pVar and *χ*^2^) for five runs starting from different random initializations of the **J** matrix. (Bottom left) Example **J** matrix for one run. (Right) Neural activity from recorded neurons (Data) and the Model RNN units. **(b)** Data presented as in Panel a for P56. **(c)** Data presented as in Panel a for P57. **(d)** Data presented as in Panel a for P58. (e) Mahalanobis distance from rest for all sixteen source currents on the familiar stimuli trials (magenta) and novel stimuli trials (cyan). Lines show mean and standard error across all five participants. Black dot indicates time of stimulus presentation. Top row is reproduced from Figure 6h.

**Figure S9.**
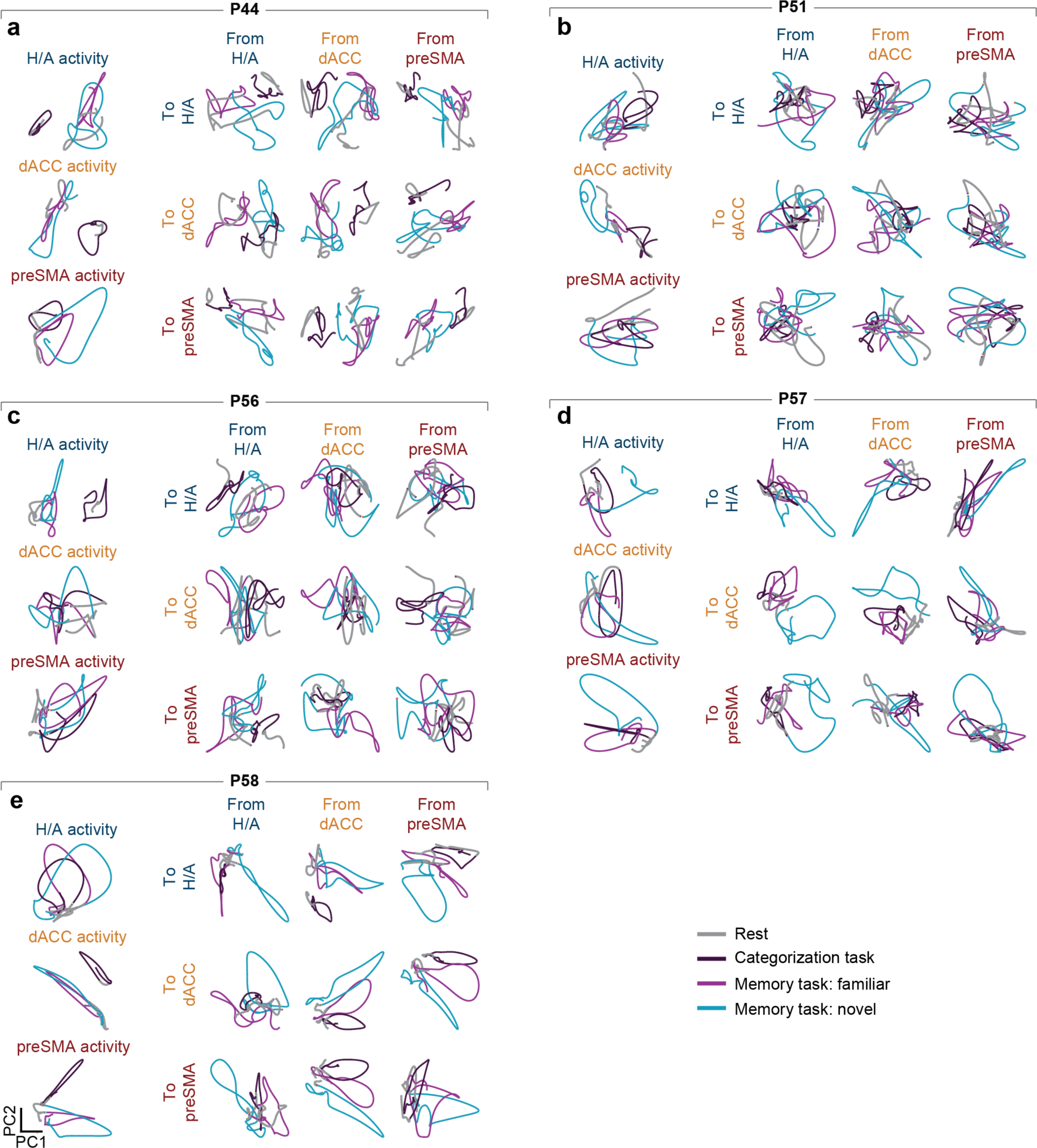
Comparison of current dynamics across tasks for the multi-region human electrophysiology dataset. **(a)** Neural and current dynamics for both tasks in P44. Each subplot shows the first two PCs of the full population activity of the three regions as well as the nine source currents during the categorization task (purple) and novel (cyan) and familiar (magenta) stimuli during the memory task. Gray shows activity at rest before the stimuli. **(b)** Neural and current dynamics for both tasks in P51. **(c)** Neural and current dynamics for both tasks in P56. **(d)** Neural and current dynamics for both tasks in P57. **(e)** Neural and current dynamics for both tasks in P58.

